# A Dynamically Regulated Designed IL-2 for Tumor-localized Signaling

**DOI:** 10.64898/2026.09.18.752703

**Authors:** Howell F. Moffett, Brian D. Weitzner, Michelle M. Lissner, Thaddeus M. Davenport, Kevin G. Haworth, Erika T. Hayes, Robert A. Langan, Leah J. Tait, Willimark M. Obenza, Bradley Hammerson, Robin L. Kirkpatrick, Laura E. Baker, John T. Crowl, Jared L. Hammer, Tina Tan, Vanessa R. Montoya, Jerry C. Chen, David S. Clausen, Paul J. Sample, Andrew H. Ng, Shujun Yuan, Rupesh H. Amin, Aaron E. Foster, Scott E. Boyken, Marc J. Lajoie

**Author notes:** These authors contributed equally to this work (Howell F. Moffett, Brian D. Weitzner, Michelle M. Lissner). These authors jointly supervised this work (Aaron E. Foster, Scott E. Boyken, Marc J. Lajoie).

## Abstract

Cytokines mediate cell-cell communication to coordinate immune responses and hold clinical promise as immunotherapies^1–3^. While natural cytokines are exquisitely regulated by immune cells, engineered cytokines have not yet matched this sophisticated regulation, limiting clinical efficacy and safety^4,5^. IL-2 is an important cytokine involved in antitumor response, but its production is disrupted in the immune-suppressive tumor microenvironment. Here we designed a regulated IL-2 module, OUTSMART designed IL-2 (dIL-2), that functions robustly in solid tumors and reduces systemic activity that could trigger toxicity: we computationally redesigned IL-2 to have a novel topology that increases stability, preserves native IL-2Rβγ interfaces, ablates IL-2Rα binding, and delivers IL-2 signaling to CD8α+ immune-effector cells. This designed cytokine is genetically regulated by a T cell activation-responsive promoter, allowing dynamic production from chimeric antigen receptor (CAR) T cells. Like wild type IL-2, OUTSMART dIL-2 drives T cell proliferation and enhances effector function. Unlike wild type IL-2, it preferentially stimulates CD8+ T cells and Natural Killer (NK) cells, avoids T regulatory cells, and stimulates CAR-T proliferation while preserving stemness, achieving durable tumor elimination in two solid tumor animal models. These design principles can be applied to create other dynamically-regulated cytokine systems that address mechanisms of complex disease.

## Introduction

The cytokine interleukin 2 (IL-2) is a potent immunotherapy agent that promotes T-cell expansion and effector function. High-dose recombinant IL-2 has produced complete responses in a subset of renal cell carcinoma and melanoma patients, but use has been limited by toxic side effects such as capillary leak syndrome^2,3^. The pleiotropic effects of IL-2 complicate its therapeutic potential, driven in part by differential expression of the IL-2 receptor (IL-2R) subunits, which can form intermediate-affinity IL-2Rβγ and high-affinity IL-2Rαβγ complexes. IL-2Rαβγ is expressed constitutively on immune-suppressive T regulatory (Treg) cells and on certain non-lymphocyte cells like lung endothelial tissue that can cause adverse events like pulmonary edema when activated by IL-2^6^. IL-2Rαβγ is critical for maximum IL-2 activity on effector CD4+ and CD8+ T cells, which upregulate IL-2Rα upon activation^7^; however, IL-2Rα is not maintained on other cell subtypes relevant to anti-tumor activity such as NK cells, which predominantly express IL-2Rβγ that is activated by both IL-2 and IL-15^8^. Thus, earlier efforts to engineer IL-2 to treat cancer focused on ablating IL-2Rα binding via mutations, protein design^9^, or chemical modifications^10,11^. These “not-alpha” therapeutics avoid undesired Treg activity but also decrease desired activity on effector T (Teff) cells^12^, and recent approaches aim to address this by targeting IL-2 variants to Teff surface markers like PD-1 or CD8β^13,14^; however, systemically-dosed protein therapeutics associated with high serum cytokine concentrations and lower intratumoral cytokine concentrations, have faced narrow therapeutic windows and/or reduced activity in the tumor compared to unmodified IL-2^4,5^.

Immune cells locally secrete cytokines in response to biological stimuli; we asked whether similar sense-and-respond programs could be engineered to coordinate a pro-inflammatory response in the tumor microenvironment. These cytokines could help translate the clinical success of CAR T-cell therapies^15,16^ to solid tumors, where efficacy is currently hindered by poor cell persistence and the inability to safely stimulate immunity within an immune-suppressive tumor microenvironment (TME)^17^. Incorporating IL-2 activity into engineered T cells can address these limitations by providing a critical “signal 3” to complement the CD3 and costimulatory signals encoded by the CAR, and by acting on endogenous cells to broaden the immune response and remodel the TME. However, existing approaches have been limited by the tradeoff between efficacy and safety: membrane-tethered IL-2^18^ or IL-15^19^, or approaches to engineer cytokine receptor signaling^20^, can improve proliferation and persistence of the engineered T cells but have limited capacity to stimulate endogenous immune cells, whereas constitutive cytokine secretion can lead to elevated circulating levels, posing toxicity risks comparable to systemically administered protein therapeutics. Inspired by the sophisticated regulation of natural cytokines, we engineered a conditionally active cytokine system specifically for solid tumors. Our OUTSMART designed IL-2 (dIL-2) exerts precise spatiotemporal control over IL-2 activity, safely driving potent responses in both engineered and bystander immune cells at the tumor site.

### Design and optimization of the OUTSMART dIL-2 system

We first designed an IL-2-based cytokine that lacks IL-2Rα binding, is genetically encodable for in vivo cellular expression, and incorporates the enhanced thermostability and engineerability of de novo proteins—all with minimal deviation from the wild-type (WT) IL-2 sequence (**Fig. 1**). IL-2 is a class I cytokine with four α-helices connected by long loops that contain the primary binding site for IL-2Rα (**Fig. 1a**). We hypothesized that removing the H1-H2 and H3-H4 loops would eliminate IL-2Rα binding, and that reconnecting the α-helices with short structured loops would preserve the native helical structure and the IL-2Rβγ interfaces while increasing stability. Starting with a model of IL-2, we generated designs that replaced the H1-H2 and H3-H4 loops using Rosetta^21^ and selected 38 relooped designs (R1.1-38) for experimental characterization. All designs eliminated IL-2Rα binding, and 33 of 38 preserved IL-2Rβγ signaling activity (**Extended Data Fig. 1**). Diluting supernatants containing the engineered cytokines further stratified their activity levels **(Fig. 1b)**. The variant with the highest combined expression and activity, R1.1, was subsequently optimized to stabilize the IL-2Rβγ binding conformation by extending the IL-2Rγ-interacting helices, redesigning the adjacent H2-H3 loop, and introducing two rationally designed mutations (R88N and I99L) to enhance IL-2Rβ interactions **(Fig. 1c)**. Candidate optimized designs were purified and screened for biological activity by measuring phosphorylated STAT5 (pSTAT5) on primary human T cells and binding affinity to IL-2Rβγ by biolayer interferometry. This optimization resulted in design R2.2, which has 86% of residues conserved at equivalent positions in WT IL-2, preserves IL-2Rβγ affinity similar to WT IL-2 (**Fig. 1d**), and retains IL-2’s characteristic helical bundle fold (**Fig. 1e**). In contrast to WT IL-2, design R2.2 is ∼20% smaller, has no detectable binding to IL-2Rα (**Fig. 1d**), is thermostable up to 95°C (**Fig. 1e**), and reduces pSTAT5 activity in Treg cells (CD4^+^Foxp3^+^) by ∼1,000-fold while preserving pSTAT5 activity in resting CD8^+^ T cells (**Fig. 1f**).

**Figure 1.**
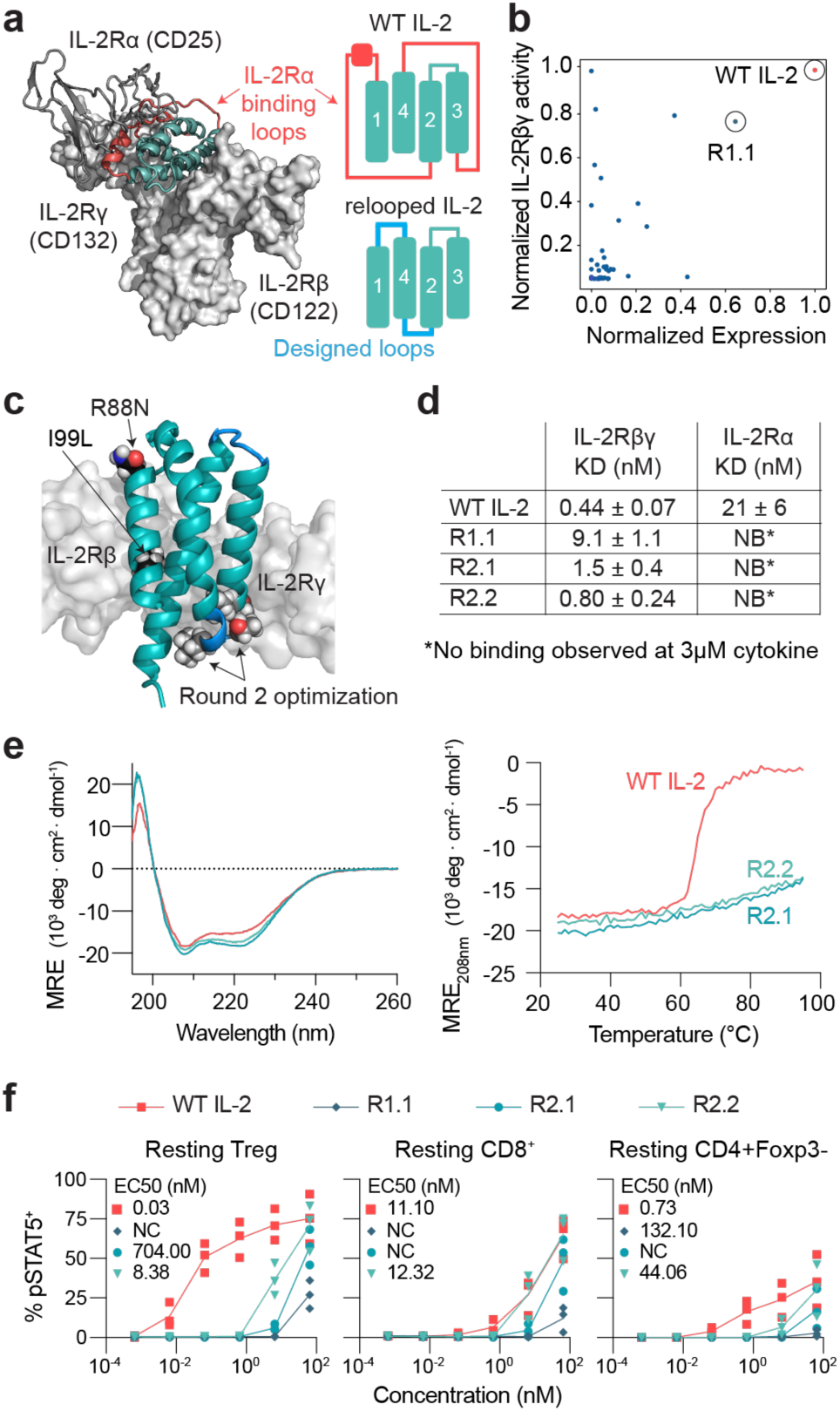
Computational design and characterization of relooped IL-2 designs that ablate IL-2Rα binding, retain IL-2Rβγ interfaces, and increase stability. **a,** Structure of WT IL-2 model, based on Protein Data Bank (PDB) ID 1M47 (cartoon representation), bound to IL-2Rαβγ from PDB ID 2B5I; IL-2Rα (CD25) shown in gray (cartoon representation), IL-2Rβ (CD122) and IL-2Rγ (CD132) shown in light gray (surface representation). Rosetta Design^21^ was used to replace the regions in red with short, structured loops (light blue) that reconnect α-helices H1-H4, generating relooped designs R1.1-38. **b,** IL-2Rβγ signaling activity (HEK-Blue reporter assay) and expression level from Expi293 supernatants diluted 1:400; designs R1.1-38 normalized to WT IL-2 levels. **c,** Optimization of design R1.1 by extending IL-2Rγ-interacting helices and redesigning the H2-H3 loop (“Round 2 optimization”) to produce design R2.1, which was further optimized by mutating two IL-2Rβ-interacting residues, R88N and I99L, to produce design R2.2. **d,** Binding affinity to IL-2Rβγ-Fc and IL-2Rα as measured by biolayer interferometry (BLI): design R2.2 binds IL-2Rβγ with a K_D_ close to that of WT IL-2 (0.80 nM and 0.44 nM, respectively), consistent with previously reported measurements for WT IL-2^35,36^. The purified proteins measured here contain STII tags; NB = no binding. **e,** Circular dichroism (CD) wavelength scan at 25℃ (*left*) and thermal melt monitoring helical signal at 208 nm (*right*). **f,** STAT5 phosphorylation (pSTAT5) in resting T cell subtypes after treatment with the indicated designed cytokines or WT IL-2. NC = not calculated.

To overcome the limitations of systemically administered ‘not-alpha’ IL-2 therapeutics, we developed a regulated system featuring two key modifications. First, the engineered IL-2 is genetically encoded for secretion from engineered T cells under the control of an activation-inducible promoter **(Fig. 2a-b)**. Second, we targeted the cytokine to CD8α to compensate for the ablated high-affinity IL-2Rα interaction, thereby enhancing specificity for endogenous T effector and NK cells **(Fig. 2c)**. To achieve this targeting, we fused variant R2.2 to a CD8α-specific VHH. The resulting construct, hereafter termed the ‘dIL-2 protein,’ bound human CD8α and IL-2Rβγ with high affinities (K_D_ = 1.27 nM and 0.53 nM, respectively), mirroring the performance of the individual components **(Extended Data Fig. 2a)**.We next measured dose-dependent pSTAT5 across different T cell populations by titrating recombinant protein. In resting Treg cells, dIL-2 protein and R2.2 were ∼10,000-fold less potent than WT IL-2 in inducing pSTAT5 activity, whereas in resting CD8+ T cells, dIL-2 was ∼1,000-fold more potent than WT IL-2 (**Fig. 2d**). In activated CD4+ T cells, dIL-2 was ∼1,000-fold less potent than WT IL-2, while in activated CD8+ T cells dIL-2 was ∼10-fold more potent than WT IL-2 (**Fig. 2d**). To assess how long-term exposure affects T cell growth and phenotype, T cells were cultured in the presence of 0.065 nM recombinant WT IL-2, dIL-2 protein, or R2.2 after 24 hours of either rest or α-CD3/28 stimulation and monitored for expansion *in vitro*. In stimulated T cells, both WT IL-2 and dIL-2 supported strong proliferation through 28 days, whereas the untargeted R2.2 was less potent (**Extended Data Fig. 2d**). While overall expansion was similar for WT IL-2 and dIL-2, the dIL-2 protein strongly biased expansion of CD8^+^ T cells (**Extended Data Fig. 2e**).

**Figure 2.**
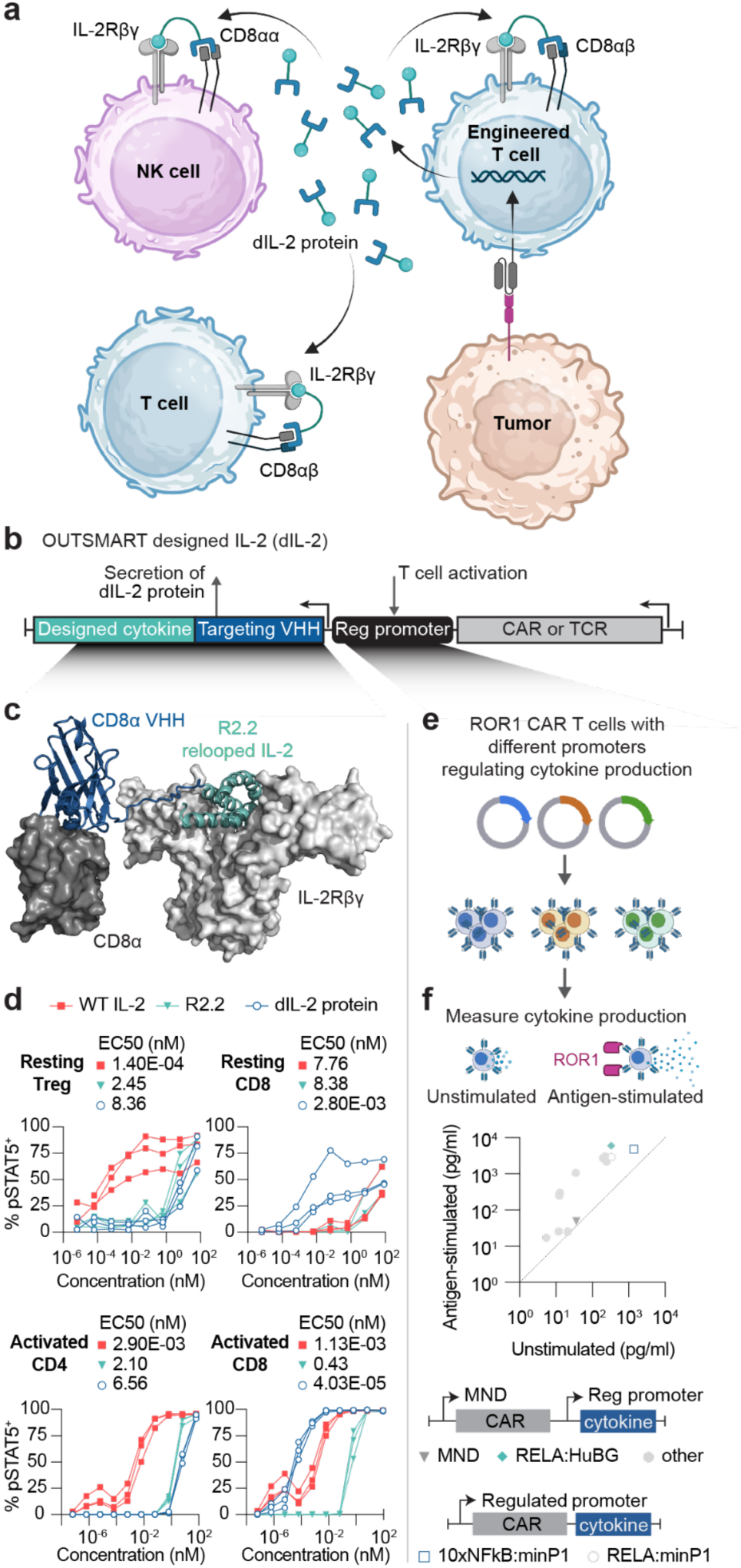
OUTSMART designed IL-2 (dIL-2). **a,** Schematic of OUTSMART dIL-2 system. In response to antigen-stimulated T cell activation, engineered T cells secrete the dIL-2 designed cytokine, which is targeted to CD8α to favor the engagement of IL-2Rβγ on endogenous NK and T cells in addition to the engineered T cells. **b,** dIL-2 is encoded in a single lentiviral vector that is broadly compatible with engineered CAR-T and TCR-T therapies. The dIL-2 protein is expressed under the control of a regulated promoter, and consists of **c,** Designed cytokine R2.2 (teal) fused to an anti-CD8α VHH (blue) via a 3xG4S linker. The VHH shown is a model for illustrative purposes (there is no crystal structure of the VHH used in this work). CD8α (PDB ID 8EW6) and IL-2Rβγ receptors (PDB ID 2B5I) are shown in surface representation. **d,** pSTAT5 activity in resting and activated T cell subtypes after treatment with recombinant protein of the indicated cytokines. **e,** Schematic of regulated promoter screen. Concatenated enhancer motifs were engineered into CAR lentiviral backbones utilizing either a dual-promoter (constitutive MND CAR with inducible cytokine) or single-promoter (CAR and cytokine expressed from the same inducible promoter) architecture. CAR-T cells were evaluated for inducible cytokine production in the presence or absence of ROR1 antigen. **f,** Regulated promoter-driven cytokine production at rest (unstimulated) versus after antigen stimulation. Illustrations in **a** were created using BioRender (https://biorender.com, Fontana, T. 2024).

We then tested the ability of dIL-2 protein to activate bystander NK and T cells, which have been shown to increase CAR-T effectiveness^22^. NK cells complement T cells in host defense and play a critical role in tumor control by integrating a range of cytokine, activation, and inhibitory signals^23,24^. A subset of NK cells express CD8α **(Fig. 3a**). Notably, these CD8^+^ NK cells exhibit enhanced cytolytic activity^25^, increased effector cytokine production^26^, and greater selectivity for MHC-I^lo^ cells, allowing them to target tumor cells that evade TIL immune pressure by downregulating surface MHC expression^27^. We measured dose-dependent pSTAT5 in purified resting NK cells by titrating recombinant dIL-2 protein. The signaling potency of the dIL-2 protein was comparable to WT IL-15 and greater than WT IL-2 in CD8+ NK cells **(Fig. 3b, left)**, whereas its potency was comparable to WT IL-2 and less than WT IL-15 in CD8-NK cells **(Fig. 3b, right)**. Despite relatively low levels of CD8 (**Fig. 3a**), limiting doses of dIL-2 protein preferentially resulted in STAT5 phosphorylation in CD8^+^ NK cells instead of CD8^-^ NK cells (**Fig. 3c**). To test NK cell proliferation and survival, we cultured CTV-labeled NK cells with and without recombinant cytokines for one week in the presence of absence of H1975 tumor spheroids; without cytokine support, NK cells cocultured with target cells did not proliferate, whereas supplementation with WT IL-2 or IL-15 caused the majority of NK cells to proliferate (**Fig. 3d**). Strikingly, dIL-2 stimulated proliferation in the majority of CD8^+^, but not CD8^-^, NK cells. We next assessed NK cell-mediated killing of H1975 tumor spheroids. NK cells without cytokine support showed no tumor killing, whereas 0.65 nM of dIL-2 protein cleared tumor spheroids almost as well as WT IL-2 and WT IL-15 (**Fig. 3e**). With ten-fold less cytokine (0.065 nM), dIL-2 performed comparably to WT IL-2 and IL-15, clearing ∼50% of target cells, whereas the untargeted R2.2 killed less well (**Fig. 3e**), highlighting the benefit of CD8-targeting when cytokine availability is limited, a likely scenario in the TME. Additionally, we evaluated whether dIL-2 would interfere with CD8 co-receptor function, which is essential for TCR recognition of peptides presented by MHC-I. We generated TCR-T cells expressing a CD8-dependent TCR targeting the tumor-associated antigen PRAME^28,29^ and activated them with peptide antigen after preincubation with supraphysiological concentrations of R2.2 or dIL-2 protein; the sensitivity and magnitude of TCR-T responses was equivalent regardless of cytokine treatment (**Extended Data Fig. 3**), indicating that the CD8α targeting of dIL-2 does not impact CD8 coreceptor function at physiologically-relevant concentrations. Taken together, these data indicate that combining a designed IL-2Rβγ agonist with an anti-CD8α VHH confers WT IL-2-like potency and enhanced cell-type specificity, enabling the support of endogenous NK cells, cancer-antigen-reactive CD8^+^ T cells, and engineered TCR-T cells.

**Figure 3.**
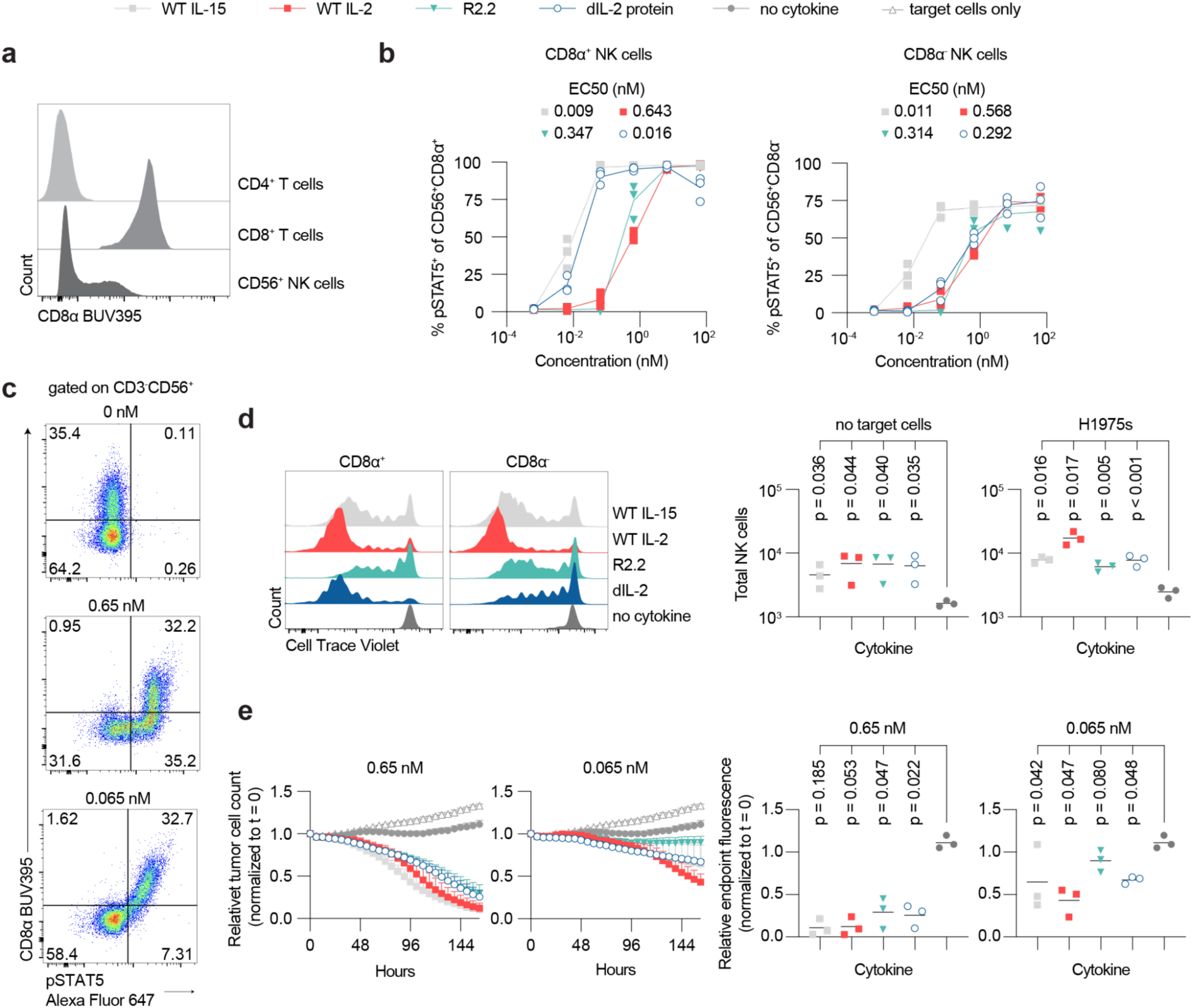
Activation of bystander NK cells. All experiments were conducted in primary NK cells isolated from 3 healthy donors. **a**, CD8a levels on CD4 T vs CD8 T vs NK cells. **b**, pSTAT5 levels induced by recombinant protein at the indicated dose of cytokine in CD3-CD56+ NK cells, split by CD8 marker expression. dIL-2 signaled more like WT IL-15 on CD8α+ NK cells and more like WT IL-2 on CD8α-NK cells. **c**, Representative flow cytometry data from one donor of pSTAT5 and CD8 expression in gated CD3-CD56+ NK cells at the indicated doses of OPR2437. **d**, NK cells were labeled with Cell Trace Violet and co-cultured at 2:1 E:T with H1975-mKate2 spheroids for 7 days, with or without supplemented recombinant cytokine at 0.65 nM. Left, Representative flow cytometry data from one donor of Cell Trace Violet expression in CD3-CD56+ NK cells split by CD8 marker expression. Right, total numbers of NK cells recovered at the assay endpoint that had been cultured in the indicated recombinant cytokines at 0.65 nM, with and without target cells; n=3 donors. **e**, NK cells were co-cultured with H1975-mKate2 tumor spheroids at 2:1 E:T alone or in combination with supplemented recombinant cytokine at 0.65 and 0.065 nM to measure NK cytotoxicity. Left, total red integrated intensity of target cells, normalized to t = 0, measured every 6 hours over one week. Right, relative endpoint fluorescence in all conditions (n=3 donors).

Lastly, to tune the regulated production in response to tumor antigen, we engineered promoters to enable stimulation-responsive expression of the designed cytokine from CAR-T cells (**Fig. 2e**). To identify CAR activation-inducible promoters, we screened concatenated enhancer motifs within two distinct lentiviral backbones. In the dual-promoter system, the CAR was driven by the MND promoter while the cytokine was controlled by the inducible promoter; conversely, in the single-promoter system, both were driven by the inducible promoter (see Methods). We identified RelA and NFkB enhancers that drove strong, simulation-dependent cytokine expression (**Fig. 2f**). Combining the inducible promoter with the CD8α-retargeted designed cytokine forms the OUTSMART dIL-2 system to achieve cell subtype specificity and tumor-localized activity.

### Enhanced CAR-T function in solid tumor models

We next evaluated the ability of OUTSMART dIL-2 to enhance CAR-T cells targeting receptor tyrosine kinase-like orphan receptor 1 (ROR1), a target for numerous hematological and solid cancers^30^. Applying the dual-promoter architecture, the constitutive MND promoter drove a ROR1-CAR and truncated CD19 (tCD19) transduction marker, while the inducible RelA promoter **(Fig. 2f)** drove either CD8α-targeted R2.2 (dIL-2), untargeted R2.2, or a WT IL-2 positive control **(Fig. 4a)**. We generated CAR-T cells from three healthy human donors and evaluated function with the ROR1-expressing H1975 non-small cell lung cancer cell line^31^. The CAR-T cells were able to induce expression of cytokine in response to ROR1 antigen (**Fig. 4b**) while maintaining secretion of endogenous IFN-γ after chronic stimulation (**Fig. 4c**). After chronic stimulation, only the OUTSMART dIL-2 and WT IL-2 CAR-T cells were able to control target cells in a spheroid model, whereas CAR-T cells lacking the cytokine module did not (**Fig. 4d**). To examine the effect of basal cytokine expression from the RelA promoter on T cell homeostasis, we tracked T cell expansion in the absence of antigen and exogenous cytokines. While CAR-T cells without the cytokine module rapidly contracted, both OUTSMART dIL-2 and WT IL-2 modules drove transient expansion followed by self-limiting contraction (**Fig. 4e**).

**Figure 4.**
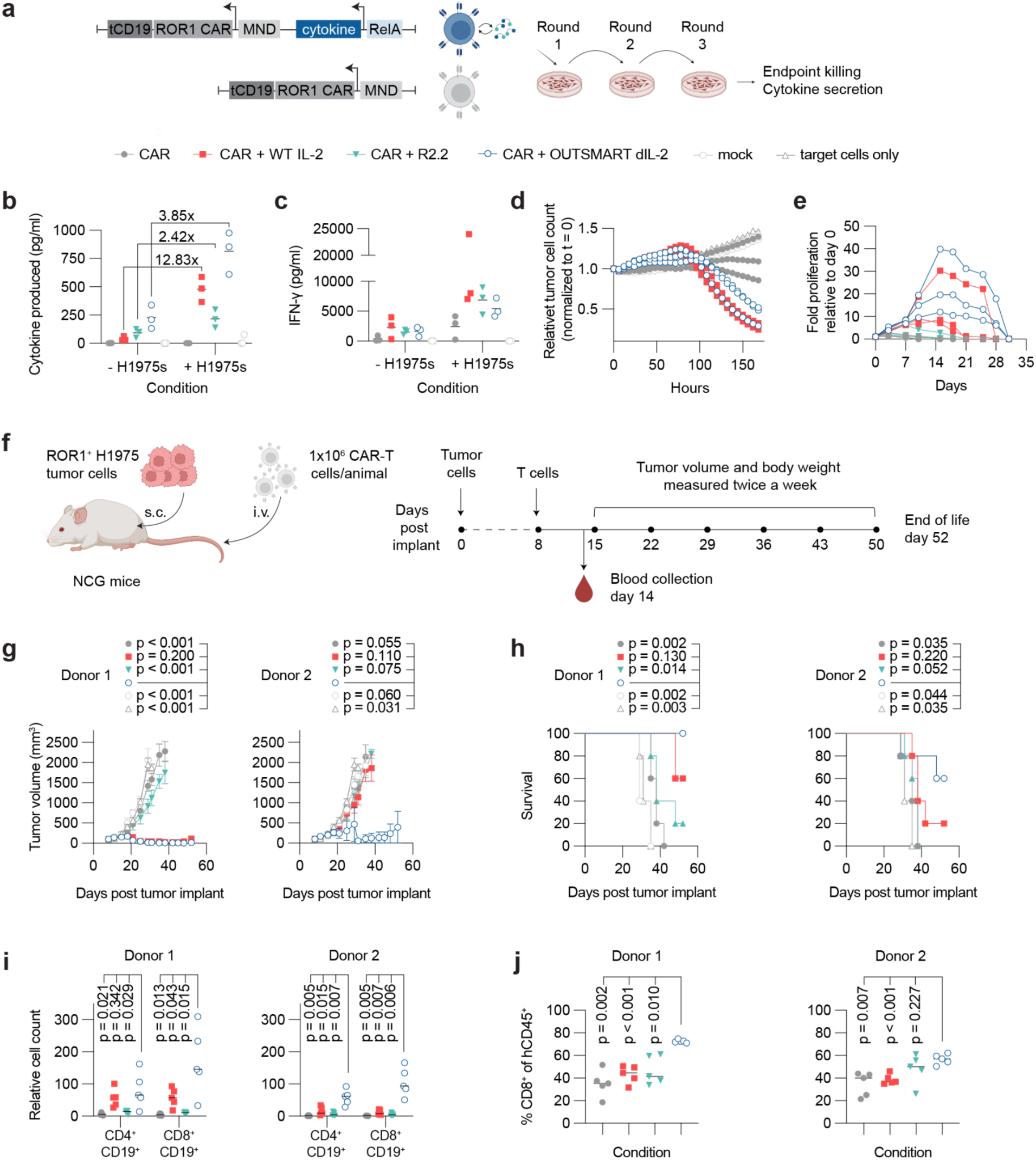
Anti-ROR1 CAR-T cells with dIL-2 demonstrate enhanced antitumor activity *in vitro* and *in vivo*. **a,** Schematic of dual-promoter lentiviral construct and experimental design for chronic stimulation experiment against three rounds with target cells followed by endpoint functional assessment. *In vitro* experiments were conducted in samples from 3 independent healthy donors. T cells transduced with the indicated constructs were cultured for 24h in the presence or absence of target H1975 cells. Here dIL-2 does not contain a GS linker, and removing the linker did not impact dIL-2 function (**Extended Data Fig. 2f**). **b-c,** cytokine production from CAR-T cells. T cells transduced with the indicated constructs were cultured +/-target H1975 cells for 24h, then supernatants were collected and analyzed by MSD assay detecting either **b,** RelA-expressed cytokine (STII-tagged) or **c,** endogenously-produced IFN-γ. **d,** Chronically-stimulated CAR-T cells were challenged with mKate2-expressing H1975 spheroids at a 1:40 ratio and assayed for target cell killing. **e,** Freshly thawed CAR-T cells were seeded in media without stimulation or exogenous cytokine. Cells were counted and split 1:2 every 3-4 days. WT IL-2 and OUTSMART dIL-2 both resulted in transient, self-limiting proliferation. **f,** Schematic illustrating a tumor xenograft study performed in immunodeficient NCG mice. **g,** Tumor volume and **h**, overall survival for mice treated with CAR-T cells from two independent healthy donors, demonstrating that OUTSMART dIL-2 outperforms WT IL-2. **i**, Counts of CD4+CD19+ and CD8+CD19+ cells for each donor from a set volume of peripheral blood collected 6 days post-CAR dose, demonstrating superior CD8+ expansion from OUTSMART dIL-2 compared to WT IL-2. **j**, Frequency of CD8+ T cells within the total human CD45+ population for each donor, demonstrating preferential CD8+ expansion for OUTSMART dIL-2. Illustrations in **a** and **f** were created using BioRender (https://biorender.com, Fontana, T. 2024).

To assess *in vivo* tumor control, we established a subcutaneous H1975 xenograft model in mice.. After 8 days, once expansion of tumor cells was apparent, mice were dosed with CAR-T cells (**Fig. 4f**). At 1x10^6^ and 2x10^6^ doses, CAR alone cells were unable to control tumor, whereas CAR-T cells with OUTSMART dIL-2 completely cleared tumors in almost all animals across both donors and doses, outperforming CAR-T cells with WT IL-2 in one donor (**Fig. 4g, Extended Data Fig. 4a**). Mice given OUTSMART dIL-2 CAR-T outperformed all other conditions at the 1x10^6^ dose (**Fig. 4h**); at the 2x10^6^ dose, OUTSMART dIL-2 mice survived comparably to the WT IL-2 condition. Six days after adoptive transfer, circulating T cell expansion mirrored expected cytokine activity, with OUTSMART dIL-2 driving the highest cell numbers, followed by WT IL-2 and R2.2 **(Fig. 4i-j, Extended Data Fig. 4c-d)**. Notably, although the targeted cytokine strongly skewed the overall population toward CD8+ T cells, it still increased the absolute number of circulating CD4+ T cells relative to CAR-T alone, indicating that the helper compartment also derives a moderate benefit **(Fig. 4i)**.

We next evaluated efficacy against a second clinically-relevant solid tumor target, mesothelin (MSLN). Applying the single-promoter architecture, 10xNF-κB:minP1 drove expression of the MSLN CAR, EGFRopt safety switch^32^, and dIL-2 (**Extended Data Fig. 5a**). Antigen exposure during a 24-hour co-culture with MSLN-expressing NCI-H1650 cells induced increased dIL-2 production **(Extended Data Fig. 5c)**. After chronic antigen stimulation, secretion of endogenous IL-2 and IFN-γ was reduced in all CAR-T cells, consistent with an exhausted phenotype, although CAR-T cells with OUTSMART dIL-2 maintained detectable IFN-γ production (**Extended Data Fig. 5d**). After chronic stimulation, only OUTSMART dIL-2 CAR-T cells were able to control MSLN+ NCI-H226^33^ spheroids (**Extended Data Fig. 5e**). We measured the expansion of CAR-T cells cultured without exogenous cytokines or further stimulation, following either CD3/CD28 activation (TransAct) or a 24-hour rest. Under these conditions, OUTSMART dIL-2 CAR-T cells exhibited a transient, self-limiting expansion profile **(Extended Data Fig. 5f-g).**

We evaluated the in vivo performance of OUTSMART dIL-2 CAR-T cells using a subcutaneous NCI-H1650 xenograft model. MND CAR-T cells controlled tumor growth at the 2x10^6^, but not 1x10^6^, dose (**Extended Data Fig. 6a**), whereas OUTSMART dIL-2 CAR-T cells maintained control at the 0.25x10^6^ cell dose (**Extended Data Fig. 6b**). Inclusion of OUTSMART dIL-2 expanded both CD4 and CD8 T cells with a strong CD8 bias (**Extended Data Fig. 6c-d**). Despite persistently high cell engraftment, circulating dIL-2 levels peaked at ∼3,500 pg/mL and then declined concurrently with tumor regression, demonstrating that OUTSMART dIL-2 expression is tumor-dependent (**Extended Data Fig. 6e**). Relative to MND CAR, OUTSMART dIL-2 improved tumor infiltration of transferred cells, nearly all of which were transduced CD8+ T cells (**Extended Data Fig. 6f-h**). Proliferation and dIL-2 production strictly required tumor antigen, as transferring OUTSMART dIL-2 CAR-T cells into tumor-free mice failed to induce cellular expansion or detectable circulating dIL-2 (**Extended Data Fig. 6i-k**). Together, these data demonstrate that OUTSMART dIL-2 expression is tumor-dependent and improves tumor control in two distinct solid tumor models *in vitro* and *in vivo*.

## Discussion

We developed OUTSMART dIL-2 to restrict cytokine activity locoregionally and cell-selectively at the tumor site, thereby avoiding systemic toxicity and Treg stimulation. To achieve this, we engineered a genetically encodable IL-2Rβγ agonist that—despite requiring only minimal mutations—adopts a more compact fold than WT IL-2 and exhibits remarkable thermostability up to 95°C. Crucially, because recapitulating a high-affinity signaling mode in the absence of IL-2Rα binding is critical for potent effector function, we retargeted this cytokine to CD8α. By coupling this targeted design with activation-inducible expression from engineered T cells, we minimized systemic exposure while establishing a robust signaling circuit that stimulates CAR-T cells, endogenous NK cells, and TILs directly at the tumor. Natural IL-2 and its receptor evolved to fight pathogens while minimizing autoimmunity^34^—regulatory safeguards that cancer readily exploits to suppress anti-tumor immunity. By combining protein design, synthetic biology, and cellular engineering, we designed a custom cytokine response tailored for cancer, wherein the cell selectivity and spatiotemporal regulation rivals that of endogenous signaling systems. This design strategy could be broadly applied to other highly potent cytokines and signaling proteins with therapeutic potential, whose native regulation is inappropriate because they evolved for fundamentally different biological purposes.

## Acknowledgements

We thank Margo Roberts and Stan Riddell for helpful discussions and feedback on the manuscript. We thank current and former members of the Outpace Bio team, including Jen Running Deer and Lesley Jones for assistance with molecular cloning, Robert Bader for assistance with *in vitro* assays, Allan Wang for assistance with lentivirus production, Erik Hermans for assistance with cloning and lentivirus production, and Maanassee Deshmukh for assistance with protein production.

## Author Contributions

M.J.L, S.E.B., B.D.W., and H.F.M. conceived of the study. B.D.W. led the computational protein design work with input from R.A.L. and S.E.B.; B.D.W., T.M.D., R.A.L., and S.E.B. performed structural analysis to optimize dIL-2 designs. M.M.L. and H.F.M. led *in vitro* functional assays and data analysis with support from E.T.H., L.J.T., W.M.O., J.T.C., T.T., and V.R.M.; E.T.H performed NK cell assays. R.L.K., J.L.H., A.H.N., J.T.C. and H.F.M. designed and performed inducible promoter screening assays. K.G.H and J.C.C. designed and performed *in vivo* experiments. B.H. performed protein production and reagent generation. L.E.B. performed binding kinetics measurements and T.M.D., S.Y., and L.E.B. analyzed the data. P.J.S. and D.S.C. assisted with visualization, data analysis, and statistics. H.F.M., M.M.L., S.E.B., and B.D.W. visualized data and generated the figures. S.E.B., H.F.M., B.D.W., and M.J.L wrote the paper, and all authors reviewed and edited the manuscript. M.J.L., S.E.B., A.F., R.A., and S.Y. supervised the work.

## AI Use Disclosure Statement

Gemini (Google) was used to assist with grammar and sentence structure in this manuscript. The author(s) reviewed and edited the output and take full responsibility for the content of the published article.

### Extended Data Figures

**Extended Data Fig. 1.**
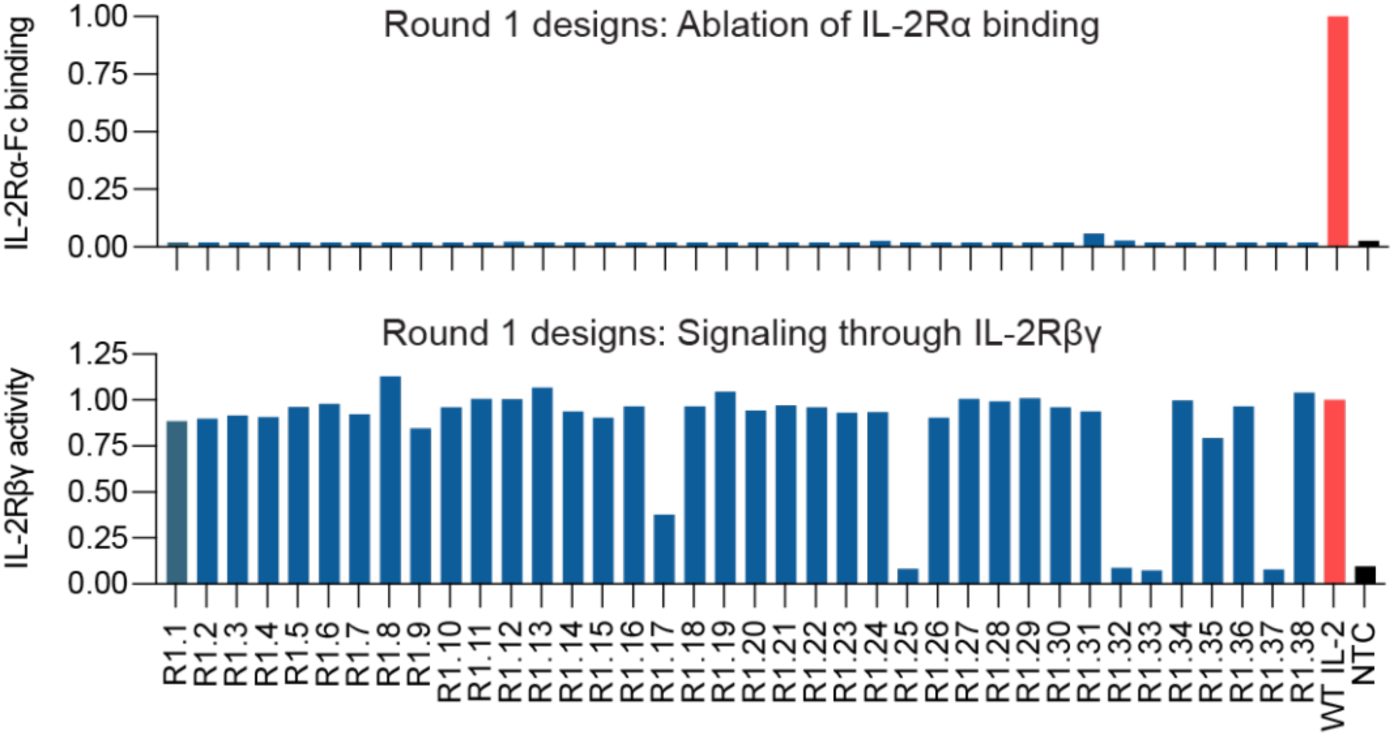
Initial screening of relooped IL-2 designs R1.1-38. Designs were expressed in Expi293 cell and supernatants collected and evaluated for binding to IL-2Rα by ELISA (*top*, normalized to WT IL-2) and IL-2Rβγ signaling activity by HEK-Blue assay (*bottom*, levels normalized to that of WT IL-2). NTC is a non-transfected control.

**Extended Data Fig. 2.**
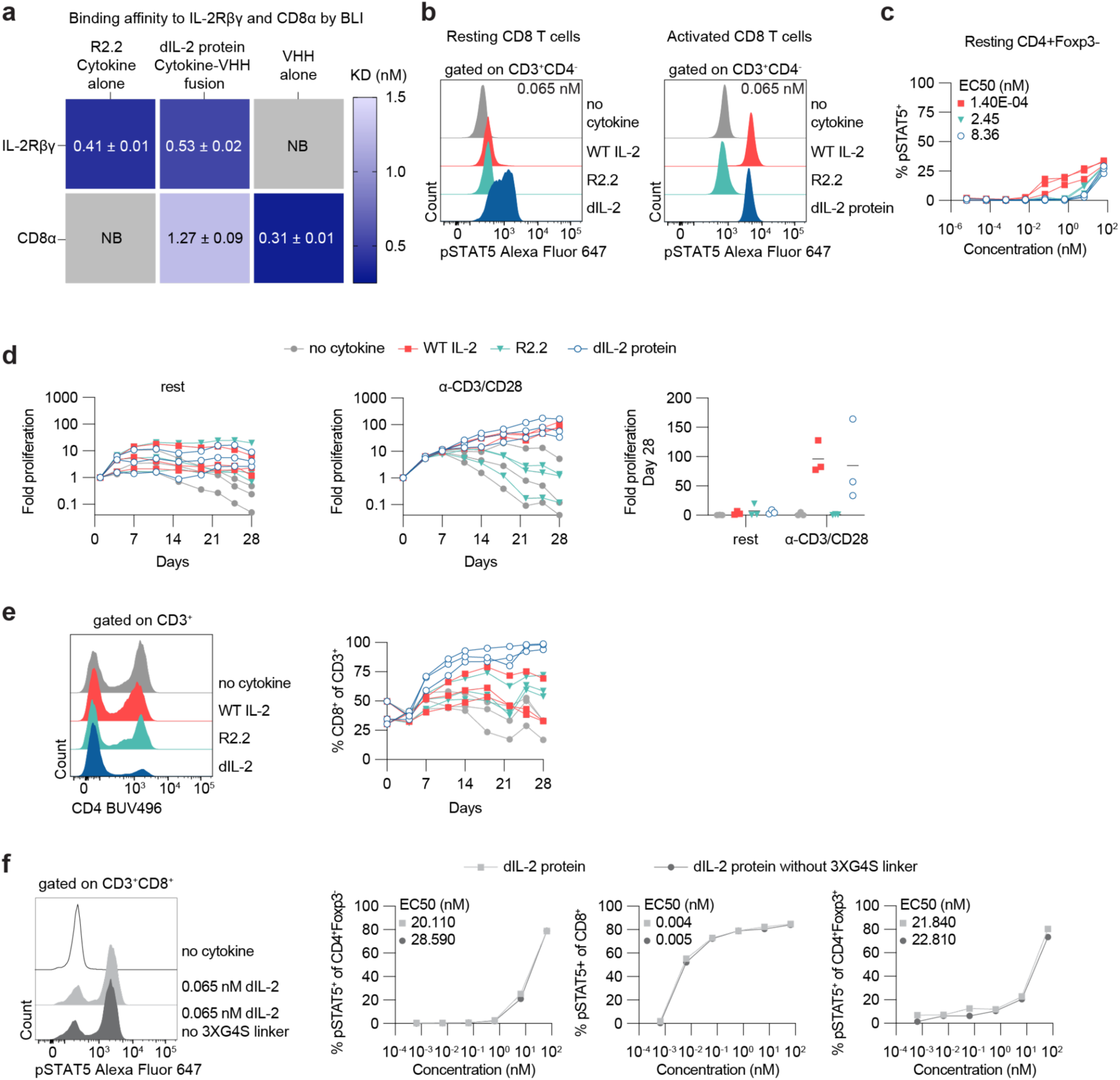
Characterization of dIL-2 protein. **a,** Binding affinity to IL-2Rβγ-Fc and CD8α measured by biolayer interferometry (BLI) for design R2.2 (untargeted), dIL-2 protein (R2.2 fused to anti-CD8α VHH), and anti-CD8α VHH alone. **b,** Representative flow cytometry plots from resting (*left*) and activated (*right*) CD8 T cells (corresponds to Fig. 2d) measuring pSTAT5 (Alexa Fluor 647) signal after treatment with the indicated cytokine at a final concentration of 0.065 nM. dIL-2 exhibits superior pSTAT5 of resting T cells and similar pSTAT5 of activated T cells compared to WT IL-2. CD8 T cells are defined by gating on CD3+CD4-signal. **c,** Compared to WT IL-2, dIL-2 exhibits less pSTAT5 in resting CD4+Foxp3-cells after treatment with the indicated cytokine. **d**, Fold proliferation of T cells that were rested or stimulated for 24h with anti-CD3/anti-CD28 (TransAct) before incubation with indicated recombinant cytokines at 0.065 nM; cells were counted and split into fresh cytokine media every 3-4 days. **e**, Frequency of CD8+ cells (defined as CD3+CD4-) in cells stimulated for 24h and grown with the indicated recombinant cytokines at 0.065 nM. By day 14, 81.3% of cells were CD8+ after addition of dIL-2 protein. **f**, GS linker does not impact dIL-2 protein function: pSTAT5 elicited by dIL-2 protein with or without 3XG4S linker between the CD8ɑ binder and cytokine elements; representative pSTAT5 staining in CD8+ T cells treated with 0.065 nM cytokine (*left*), and pSTAT5 in resting T cell subtypes after treatment with the indicated designed cytokines (*right*).

**Extended Data Fig. 3.**
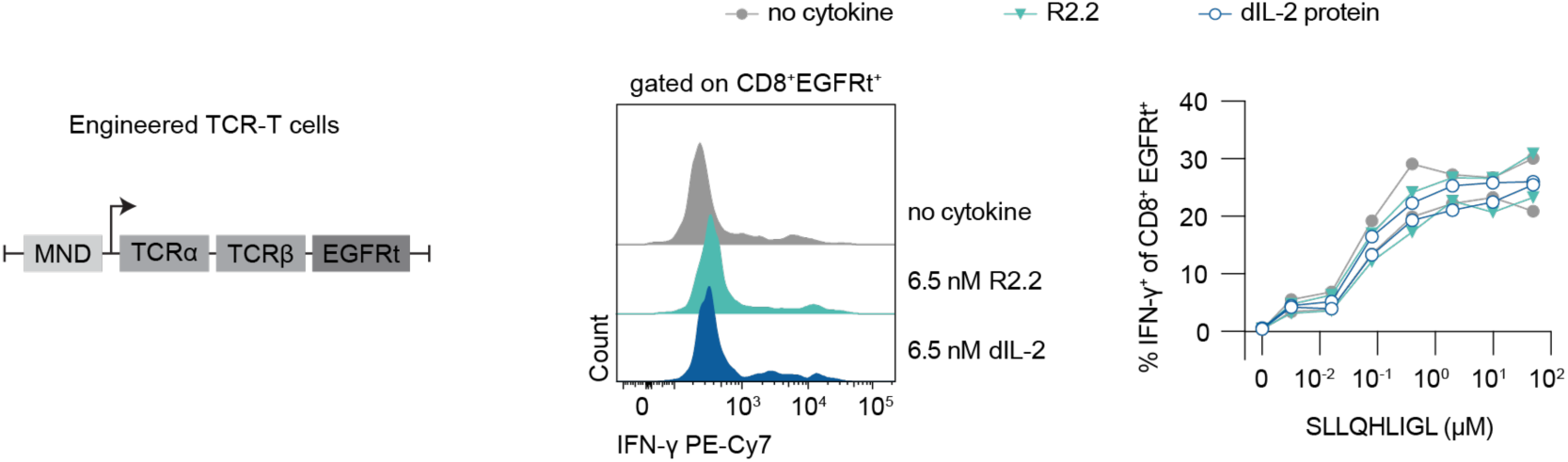
CD8 binding of dIL-2 does not impede TCR function. Engineered T cells over-expressing a CD8-dependent TCR specific for PRAME antigen SLLQHLIGL were evaluated for activation (IFN-γ) in response to SLLQHLIGL peptide in the presence or absence of R2.2 or dIL-2 protein.

**Extended Data Fig. 4.**
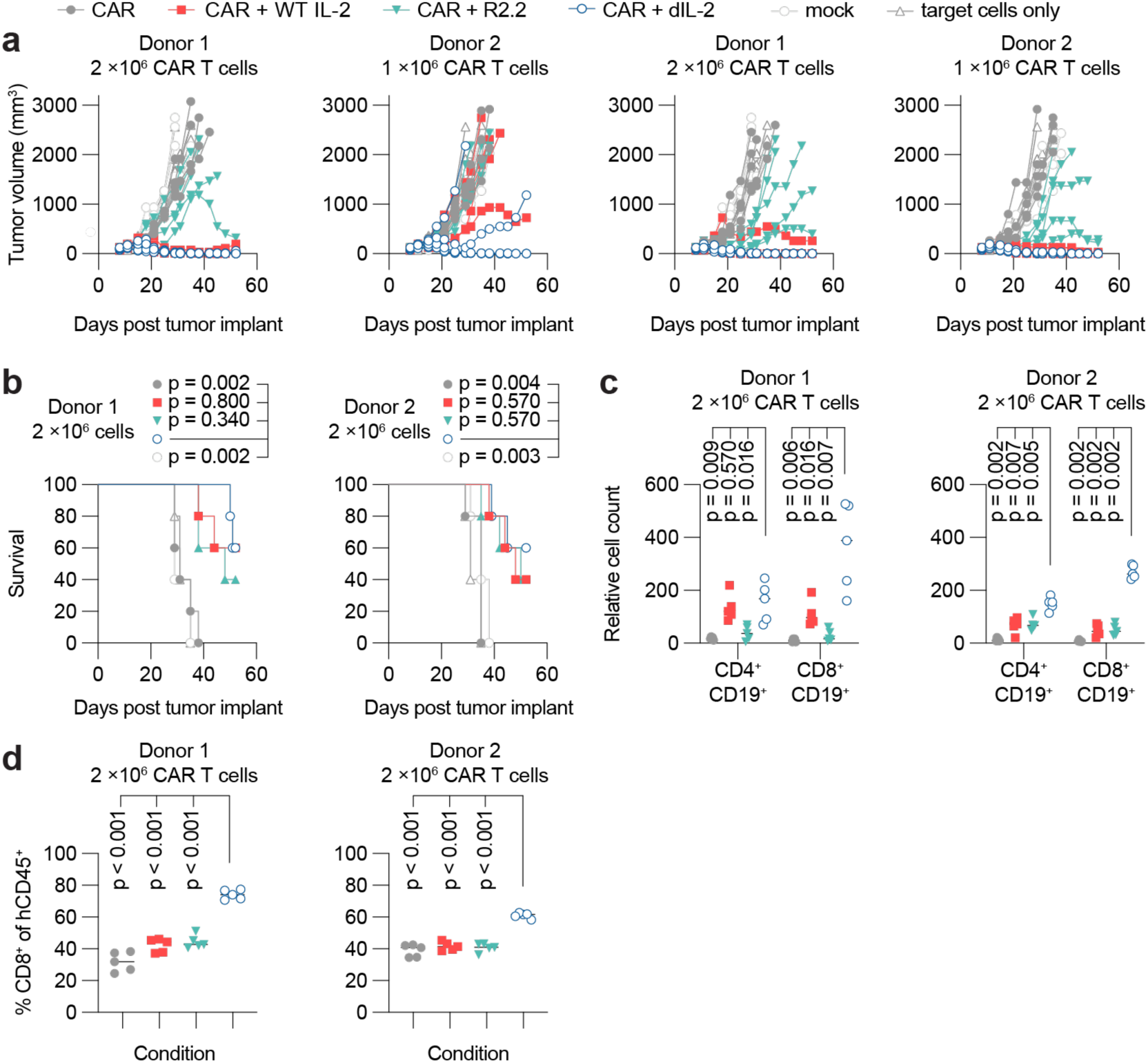
Anti-ROR1 CAR-T cells with OUTSMART dIL-2. A xenograft study was performed in immunodeficient NCG mice implanted with H1975 tumor cells (Fig. 4). **a,** Individual tumor volume plots (measured as mm^3^) for each human donor and CAR dose condition. The first data point is Day 8 post-tumor implant and corresponds to the first measurement post CAR-T cell dose. Individual lines representing individual mice stop when the animal is removed from study. CAR T cells with OUTSMART dIL-2 exhibit improved antitumor efficacy compared to CAR T cells with WT IL-2. **b,** Survival plots for each group plotted in similar order as the tumor volume graphs and each census change within a group is indicated by a corresponding symbol on the survival line graph; at the end of the study, some mice exhibited graft-versus-host disease (GVHD), which contributed to decline in survival. **c,** Cell counts of CD4^+^CD19^+^ and CD8^+^CD19^+^ for each donor and dose condition from a set volume of peripheral blood collected 6 days post-CAR dose. Compared to WT IL-2, OUTSMART dIL-2 leads to superior CD8+CD19+ T cell expansion. **d,** Frequency of CD8^+^ T cells within the total human CD45^+^ population for each donor and dose condition. Legends for each series of graphs are shown on the top of the figure.

**Extended Data Fig. 5.**
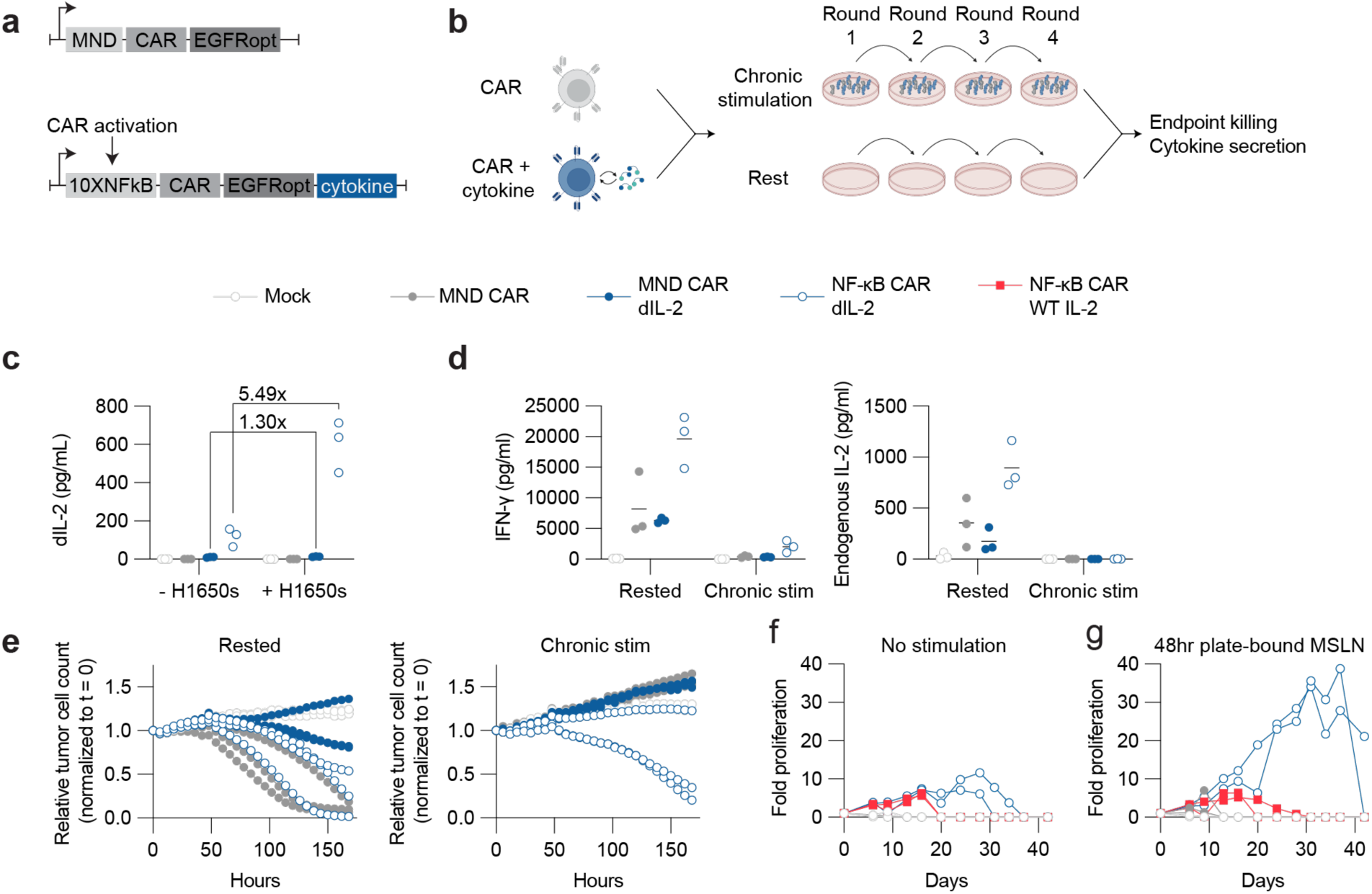
Anti-MSLN CAR-T cells with OUTSMART dIL-2 demonstrate enhanced antitumor activity *in vitro*. **a,** Schematic of the single promoter lentiviral constructs **b,** Experimental design for chronic stimulation experiment of CAR-T cells for 4 rounds with plate bound antigen over 10 days followed by endpoint functional assessment.**c,** Supernatants from freshly thawed CAR-T cells cocultured +/-H1650 target cells for 24h were assessed for OUTSMART dIL-2 production by MSD. **d,** Supernatants from chronically-stimulated or rested CAR-T cells incubated with H1650 target cells for 24h were assessed for IFN-γ and endogenous IL-2 production by MSD. **e,** Chronically-stimulated or rested CAR-T cells were cocultured with H226 target cell spheroids at a 1:20 E:T and killing was measured using the Incucyte platform. Freshly thawed CAR-T cells were seeded **f,** in media without stimulation or exogenous cytokine or **g,** stimulated with plate-bound antigen for 48h, then seeded in media without stimulation or exogenous cytokine. Cells were counted and split 1:2 every 3-4 days.

**Extended Data Fig. 6.**
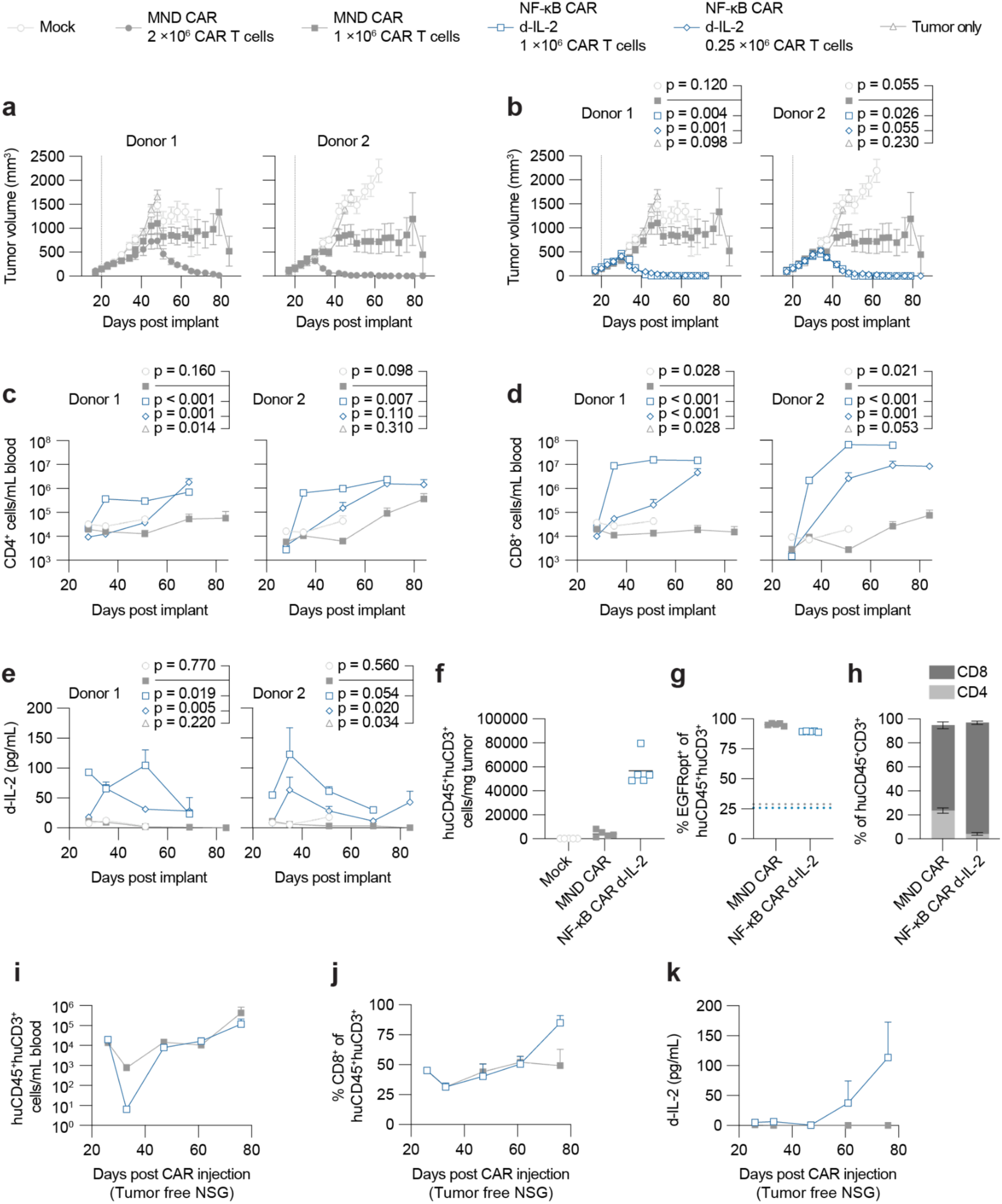
Anti-MSLN CAR-T cells with OUTSMART dIL-2 demonstrate enhanced antitumor activity *in vivo*. **a-h**, Xenograft studies performed in immunodeficient NSG mice engrafted with human MSLN+ H1650 tumor cells were treated with either MND CAR only or OUTSMART dIL-2 CAR-T cells with both CAR and dIL-2 are under control of a 10xNF-κB:minP1 promoter. **a,** Mean tumor volume in mice treated with 1x10^6^ or 2x10^6^ MND CAR cells. **b,** Comparison of tumor volume in mice treated with 1x10^6^ MND CAR versus 1x10^6^ or 0.25x10^6^ OUTSMART dIL-2 cells. **c,** CD4+ T cell counts per mL of blood. **d,** CD8+ T cell counts per mL of blood. **e,** Serum dIL-2 levels over time from animals presented in **b**. **f,** Tumor infiltrating T cells in mice treated with MND CAR or OUTSMART dIL-2 CAR T cells, assessed 15 days after CAR-T injection. OUTSMART dIL-2 dramatically increases intratumoral T cell counts. **g,** percentage of tumor infiltrating cells which express the EGFRopt CAR transduction marker, with horizontal dashed lines representing input percentage. **h,** CD4+ and CD8+ tumor infiltrating cell frequency between MND CAR and OUTSMART dIL-2 CAR. (**i-k**) Pharmacodynamics of MND CAR or OUTSMART dIL-2 CAR in the absence of antigen in tumor-free NSG mice, demonstrating that proliferation is antigen dependent (late expansion is likely due to alloreactivity given that the timing of expansion is as-expected for the onset of GVHD in this model). **i,** Circulating huCD45+ huCD3+ cells. **j,** Circulating huCD45+ huCD3+ CD8+ T cells. **k,** Serum dIL-2 over time in the blood of mice treated with 1x10^6^ CAR T cells. Error bars represent standard error of the mean (SEM), vertical dotted lines represent CAR dose. Mean lines stop being plotted with >40% of mice are off study within a group.

## Methods

### Key reagents and materials used in this study

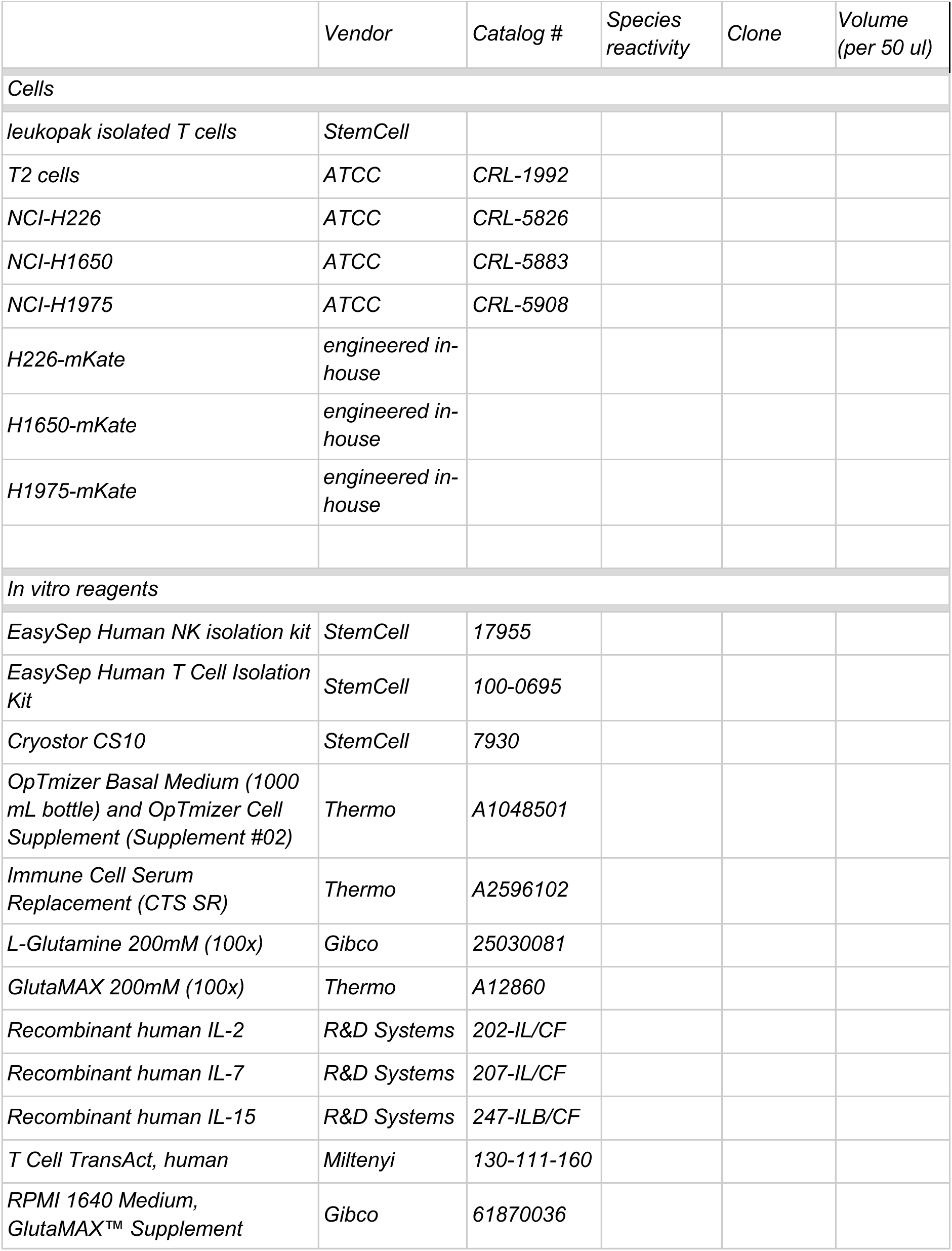
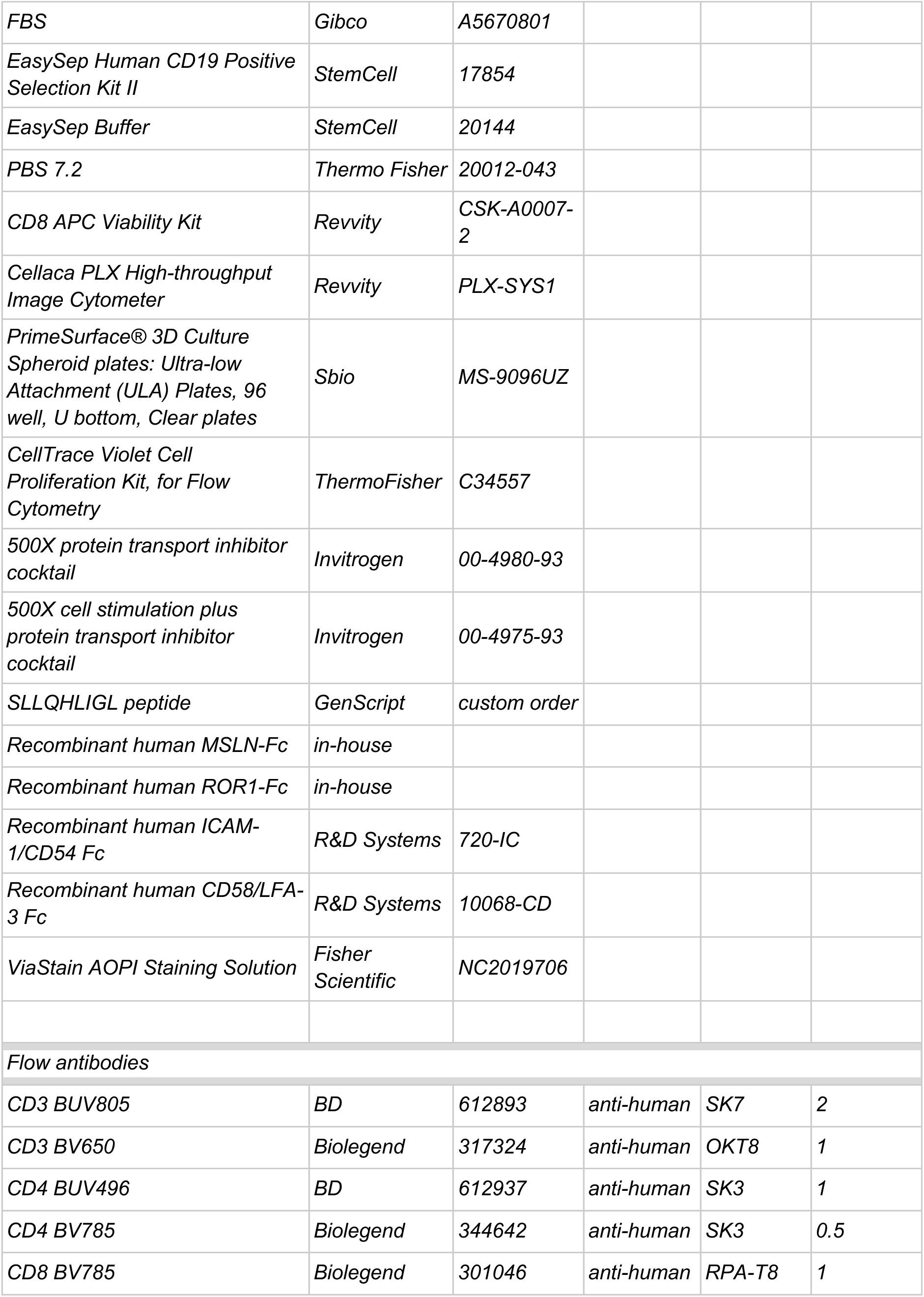
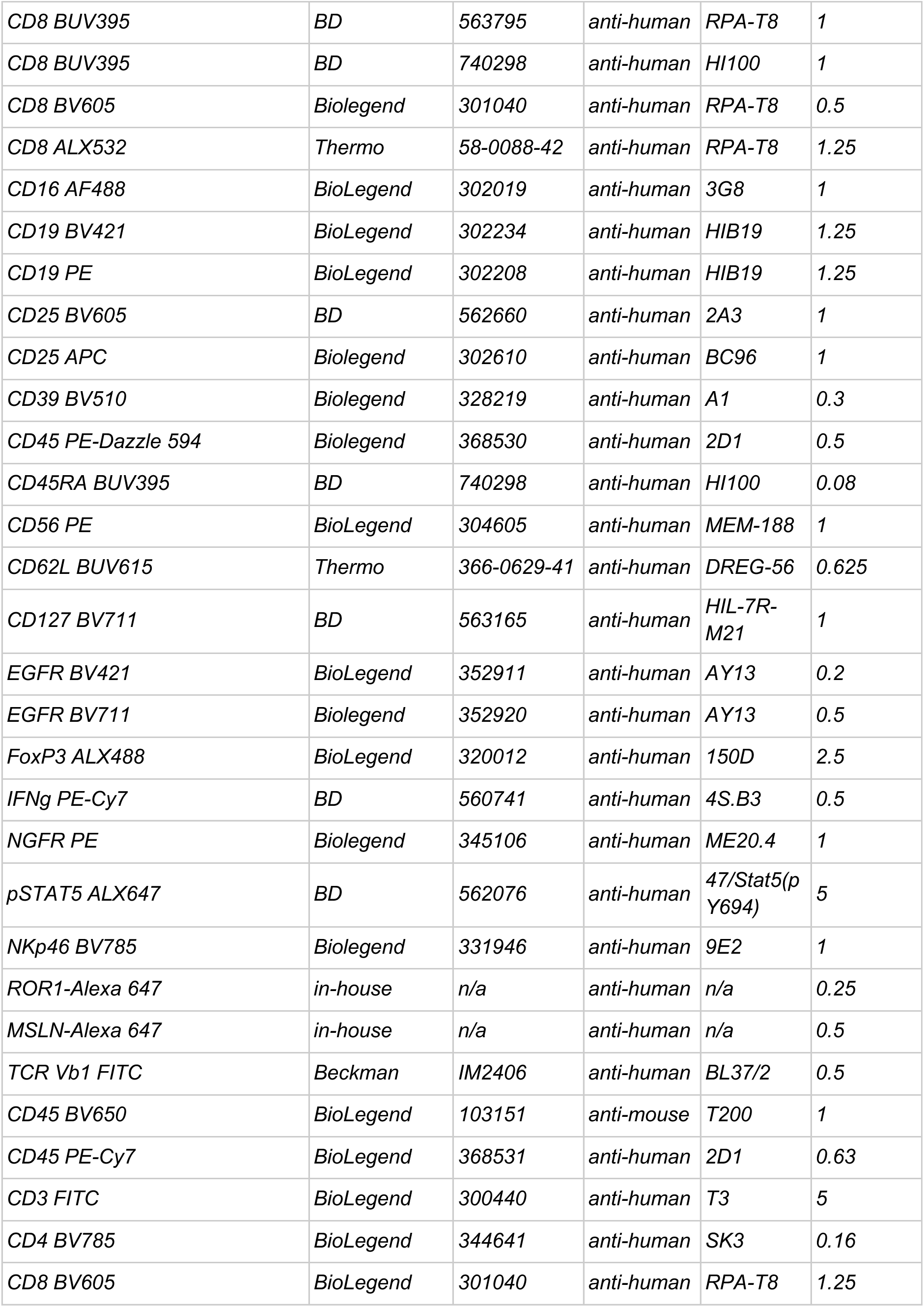
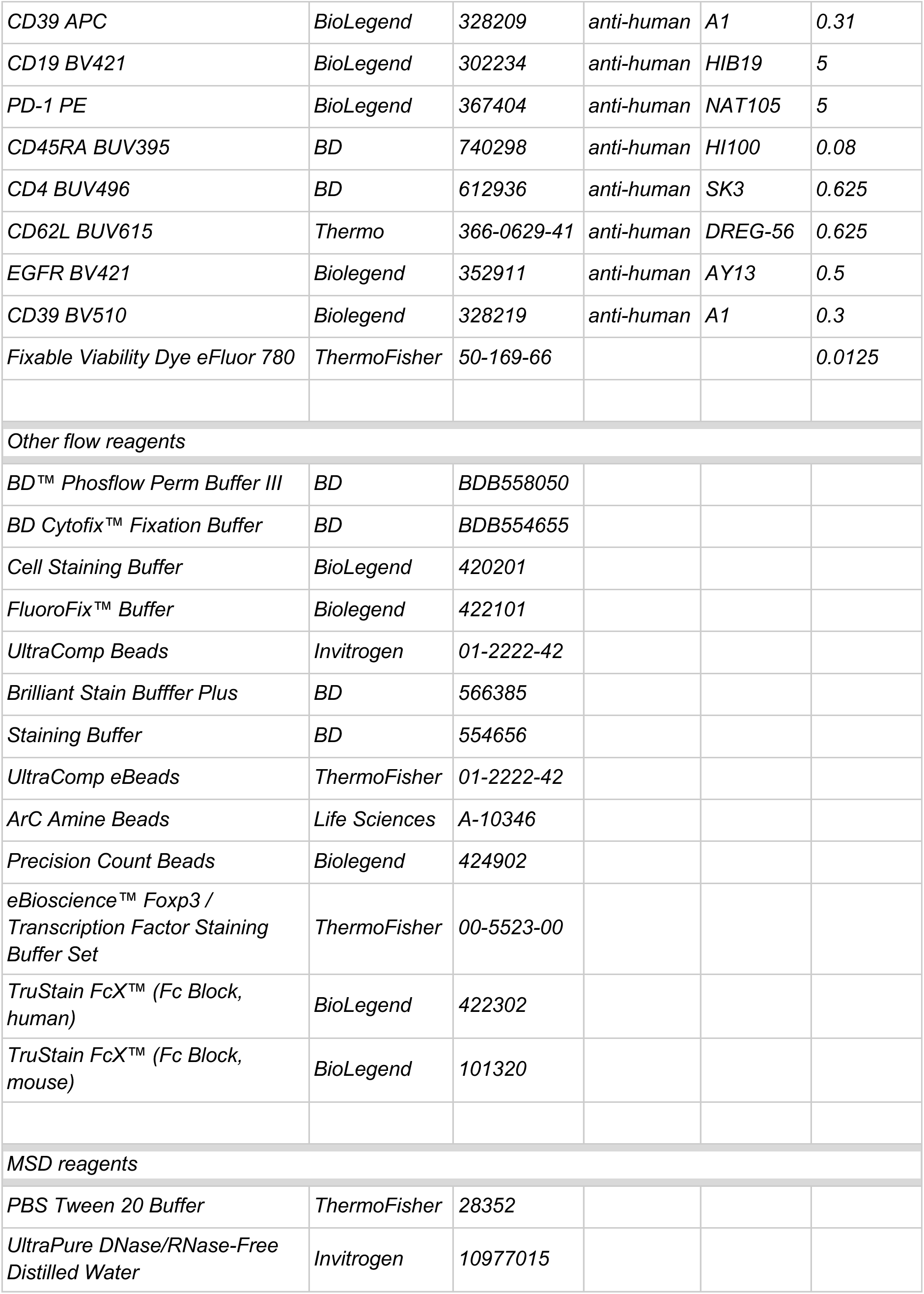
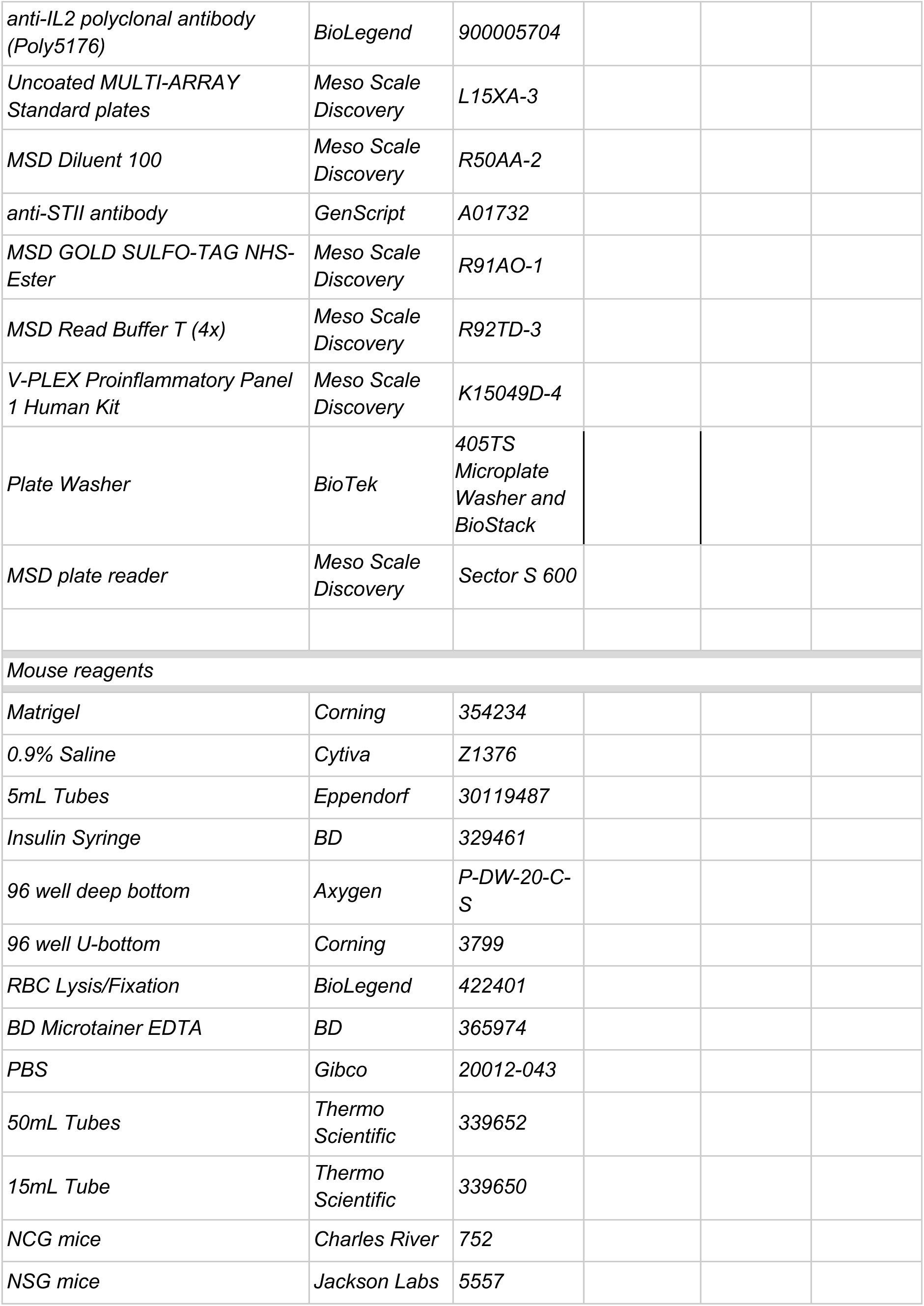
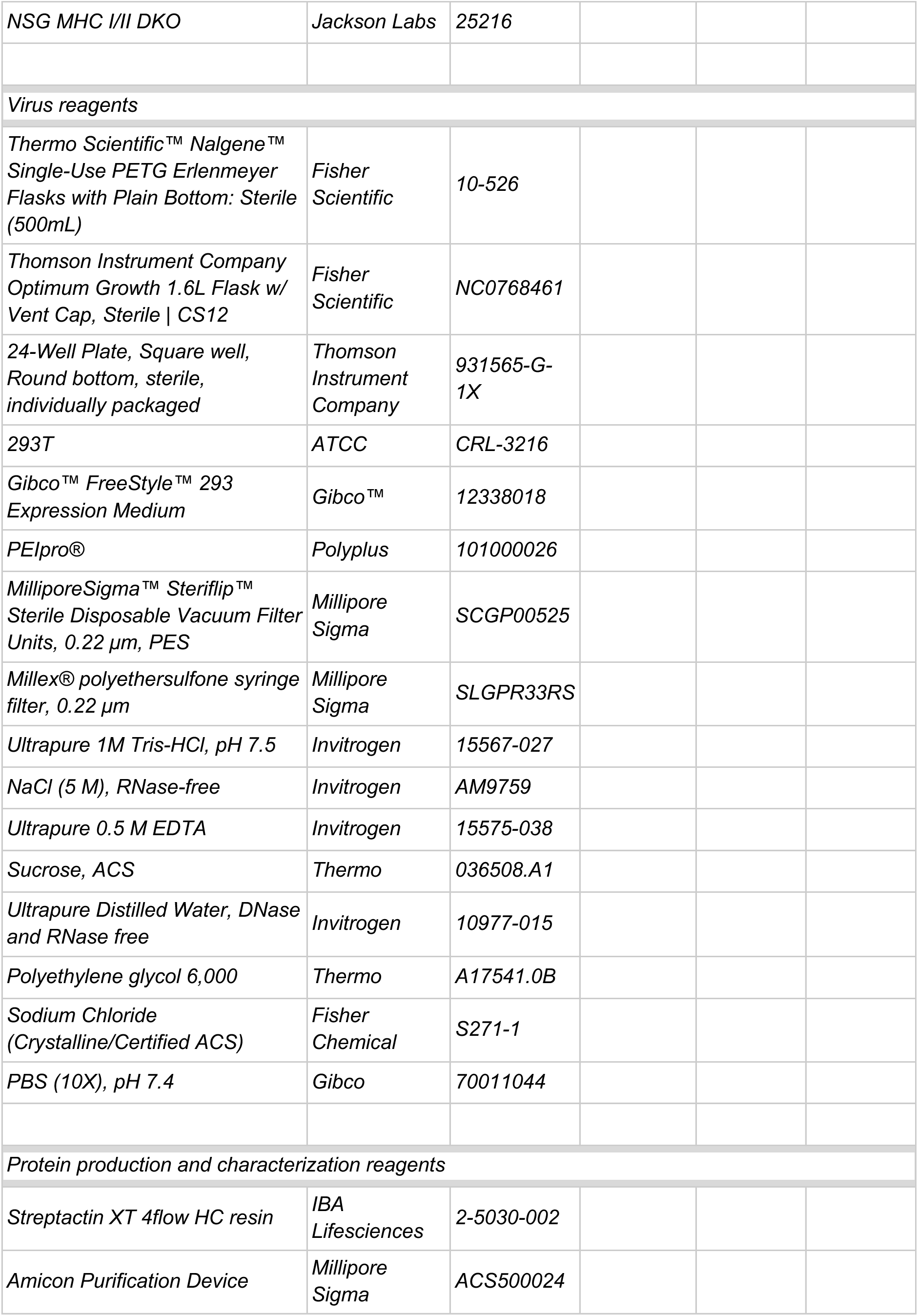
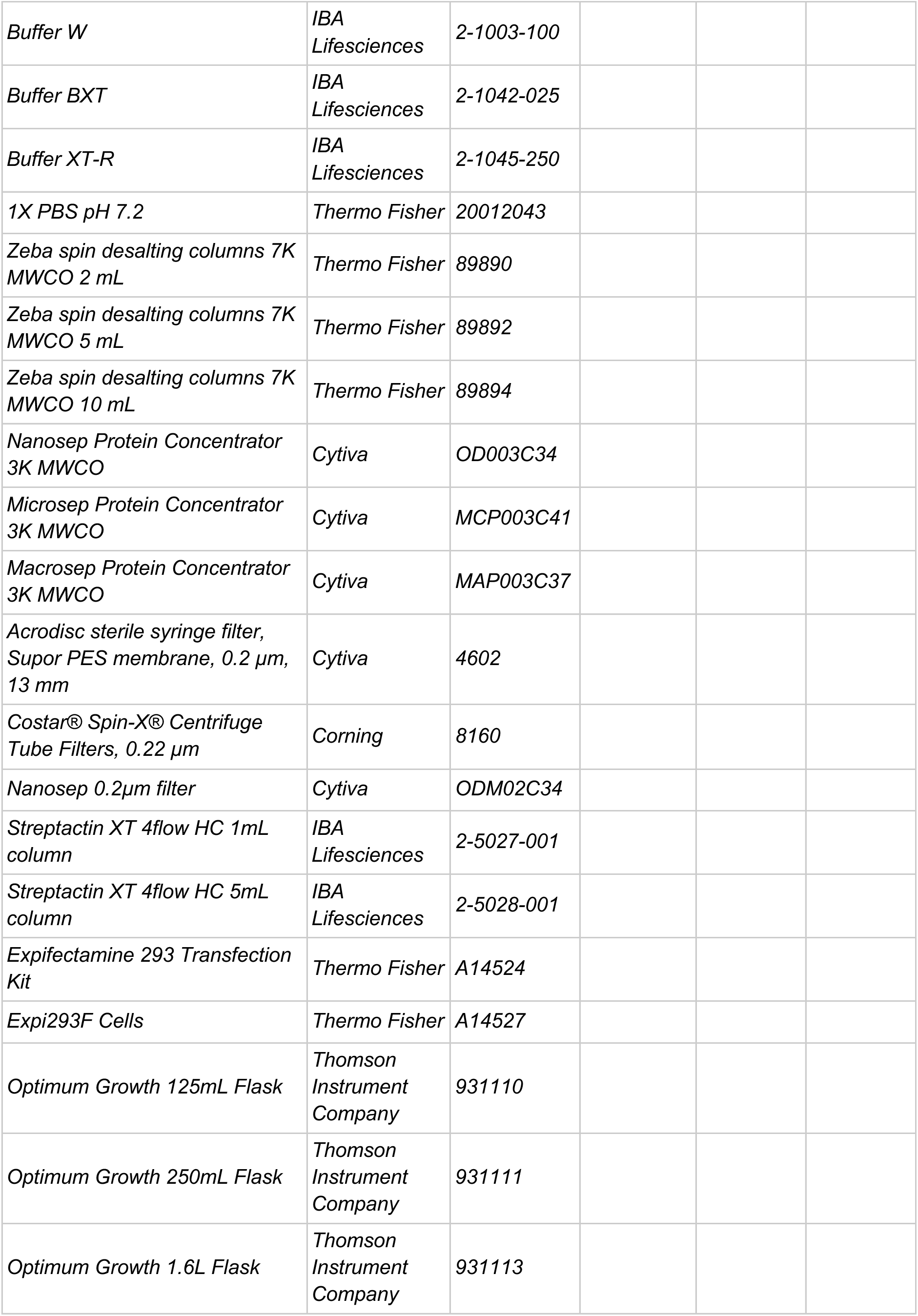
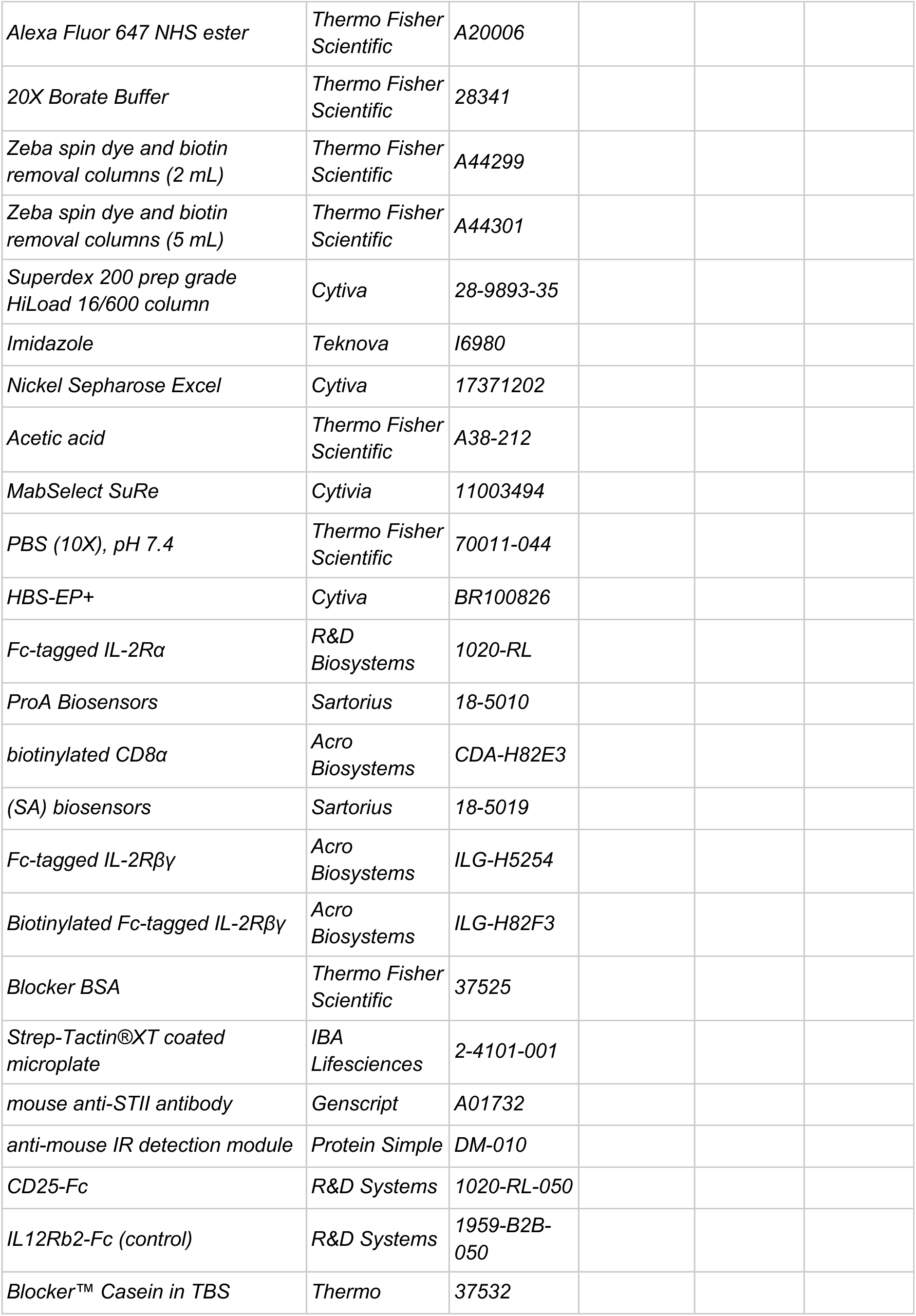
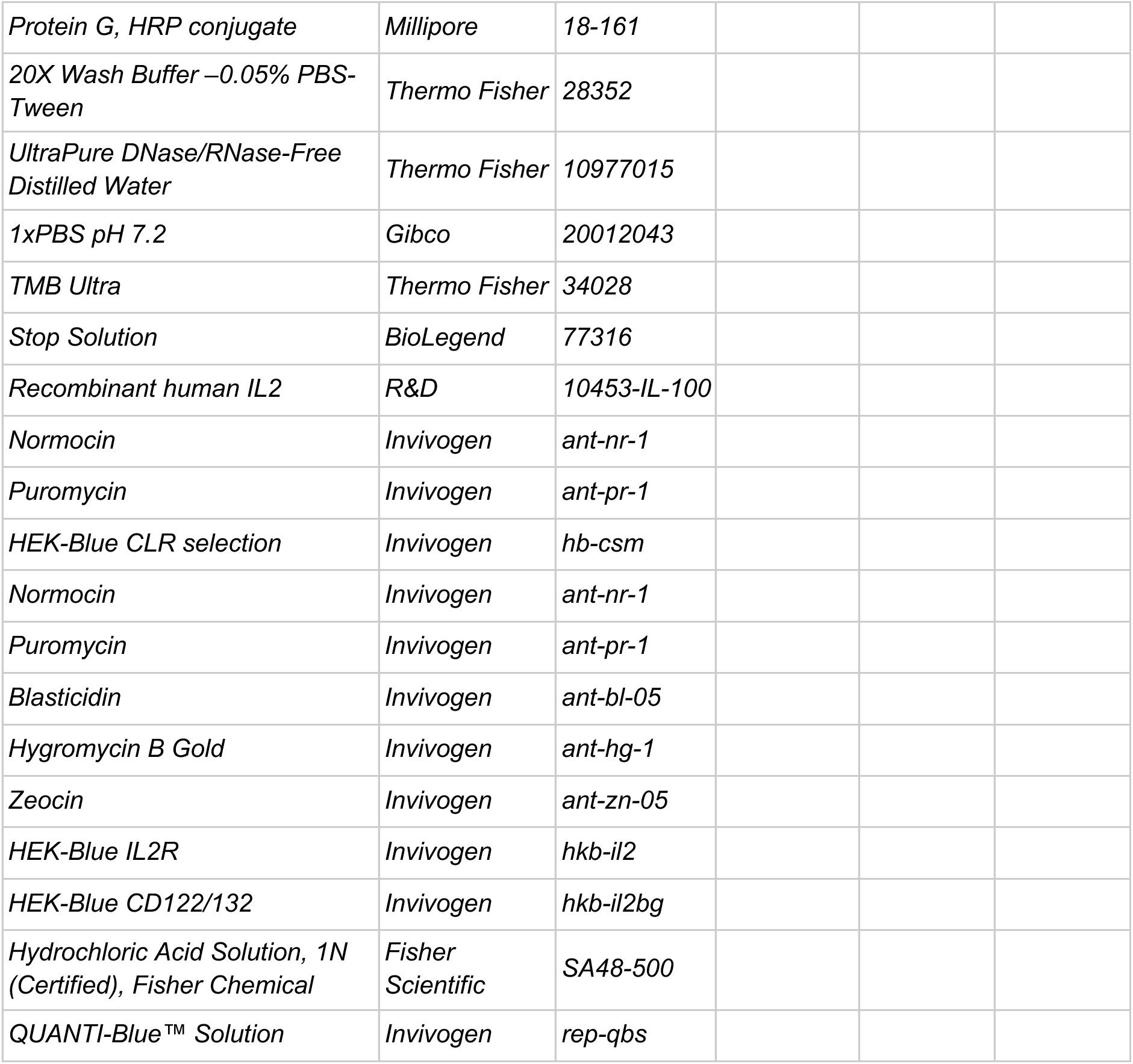

### Protein Design

#### Backbone and Sequence Design

The IL-2 structural model that was used as a starting point for design is Protein Data Bank (PDB) ID 1M47, which was downloaded from RCSB (https://www.rcsb.org/structure/1m47) and prepared for Rosetta Design^21^ using a constrained relax protocol which restrains the backbone atoms to within a net 0.5 Å deviation from the crystal structure. This serves to maintain the backbone structure that was experimentally determined while equilibrating it to the Rosetta energy function. Missing residues from the crystal structure were modeled with Rosetta Remodel^37^ using a blueprint file to define the existence and identity of the missing residues that needed to be added back.

To initiate backbone design, we manually adjusted the order of helices in the text PDB file to H1, H4, H2, H3 while maintaining only the H2-H3 structured loop. Between H1 and H2, in some design instances we removed a single turn of the helix from H1, and in others we kept that turn intact. Backbone design between H1-H4 and H4-H2 was performed in Rosetta using the NearNativeLoopCloser and DirectSegmentLookup movers. We allowed these algorithms to remove between 0-4 residues, inclusive, on each side of the loop to find matches in a structural database of helix-turn-helix motifs. This database contains helix-loop-helix motifs from the PDB with 50% or less sequence identity with loop conformations (described by a coarse-grained Ramachandran description with A corresponding to α-helical conformations; B to β-strands; G to left-handed helices; E to the remaining conformational space; and O for residues with cis-peptide bonds) limited to "GB", "BB", "GBB", "BAB", "BBB", "GBBB", "GABB", and "BBBB"). The NearNativeLoopCloser and DirectSegmentLookup movers use similar algorithms to match the RMSD of motifs in the database to the desired connection points on the helices in our cytokine and selecting any match within a 0.4Å cutoff. The Rosetta PackRotamersMover performed a fixed-backbone, monte-carlo simulated annealing sequence design algorithm on each motif added from the database, and residues within 5Å of the newly placed residues. The backbone of the resulting loops were minimized in the Rosetta ref_2015 cartesian energy function using a lbfgs_armijo_nonmonotone minimization algorithm and any loops with a chainbreak, defined as a backbone bond length with deviation at least 0.13Å from ideal 1.33Å.

#### Design Selection Criteria and Filtering

Since two loops were designed independently of each other, every combination of H1-H4 and H4-H2 loop was considered for final selection and subjected to a battery of computational assessments for overall design quality. The most important metrics we used were length of the design(between 65 and 105 amino acids total), 2 or fewer unsatisfied hydrogen-bonds based on the Rosetta UnsatHBondsFilter, a computed isoelectric point of less than 6.4, Rosetta rama_prepro score of -10 REU or less (a metric of backbone quality conditioned on sequence), and a total Rosetta score of less than -100 REU.

To further improve the designs, we ensured the presence of known α-helical characteristics: amino acid identity at the helical N and C terminal capping residues^38,39^ and overall composition of charged residues to complement the helical dipole. Visual inspection was then used both to filter out structures with helices that deviate from their positions in wild-type IL-2 or with loop conformations that appeared physically unrealistic and to identify designed positions with backbone atoms that aligned with those of WT residues that were removed for the design process. Residues at these positions were mutated to the WT identity. This filtering process resulted in 38 designs that met the above criteria.

Structure visualizations and figures were created with PyMOL v.2.5 (https://github.com/schrodinger/pymol-open-source, Schrodinger)

#### Experimental screening of Relooped IL-2 Designs

The initial screening of STII-tagged, relooped backbones was performed by transfecting Expi293 cells in a 96 well deep-well block. Five days post-transfection, the block was centrifuged at 4000xg for 10 min and supernatants were transferred to 96 well plates for downstream analysis, including IL-2Rα-Fc (CD25-Fc) ELISA, HEK-Blue IL-2R activation, and Jess western to measure expression.

IL-2Rα-Fc (CD25-Fc) ELISA was performed by capturing STII-tagged proteins out of undiluted supernatants on Streptactin XT coated plates (IBA Lifesciences) by incubation for 1 hour at room temperature. Plates were then washed with PBS-T (Thermo), and incubated with the primary detection reagent, CD25-Fc (R&D Systems), diluted to 0.5 μg/mL in dilution buffer (50:50 casein block buffer

(Thermo):PBS-T) for 1 hour at room temperature. Plates were again washed with PBS-T, and then incubated for 45 minutes with the secondary detection reagent, protein G-HRP (Millipore) at 1:5,000 in dilution buffer. Plates were washed again prior to the addition of TMB-Ultra (Thermo) and 2M HCl (Fisher Scientific) to stop development. Absorbance at 450 nm was measured on a Synergy H1 plate reader (Agilent).

The activity of relooped IL-2 molecules in Expi293 supernatants (undiluted, and diluted 1:10, 1:50, 1:100, 1:200, and 1:400) was assayed on HEK-Blue IL2R cells (Invivogen) according to the manufacturer’s instructions. After an overnight incubation of 18-20 hours, 20 μL of each HEK-Blue IL2R supernatant was added to 180 μL of Quantiblue (Invivogen). The Quantiblue reaction was developed for 30 min at 37℃ prior to measuring absorbance at 640 nm on a Synergy H1 plate reader.

Expression of relooped IL-2 molecules in Expi293 supernatants was evaluated using a Jess capillary electrophoresis western blot apparatus (Protein Simple). Samples were prepared under reducing conditions according to the manufacturer’s instructions, separated on a 25 well 2-40 kDa capillary cartridge, and probed using a primary mouse anti-STII antibody (Genscript) diluted 1:100 and an anti-mouse IR secondary antibody detection module (Protein Simple). The relative expression of relooped IL-2 molecules compared to STII-tagged WT IL-2 was calculated using integrated fluorescence intensity values for peaks of the expected molecular weights.

### Protein production and biophysical characterization

#### Recombinant protein production and purification

Proteins were produced by transient transfection using the Expi293 Expression system (Thermo Fisher) and plasmids encoding the following proteins tagged with an N-terminal human Igκ signal peptide under the control of the CMV promoter: IL-2 variants with a C-terminal Strep-II (STII) tag; MSLN isoform 2 (residues 296-582) with an Avi-His tag; or human ROR1 extracellular domain bearing a c-terminal human IgG1 Fc domain. Five days post-transfection, supernatants were harvested by spinning for 10 min at 4,000 x g at 4°C, filtered with 0.2µm PES syringe or vacuum filters (Cytivia/Corning), and held at 4°C until purification.

Batch purifications were done using Amicon purification devices and a table top centrifuge. Buffer exchange was done into 1X PBS pH 7.2 and samples were sterile filtered. STII-tagged proteins were purified with Streptactin XT resin and manufacturer-recommended buffers (IBA Lifesciences). His-tagged MSLN was purified with Nickel Sepharose Excel (Cytiva) in PBS at pH 7.4 with 0.5M NaCl and eluted with 0.5M Imidazole.

FPLC purifications were done using an AKTA Pure for affinity chromatography, followed by preparative size exclusion chromatography in 1X PBS pH 7.2 using a Superdex 200 Column (Cytiva) and sterile filtration. Affinity chromatography for STII-tagged proteins was performed with Streptactin XT columns and manufacturer-recommended buffers (IBA Lifesciences) while affinity chromatography for His-tagged MSLN proteins was performed with His-Trap Excel (Cytiva) in PBS at pH 7.4 with 0.5M NaCl and eluted with 0.5M Imidazole in a linear gradient. Affinity purification of ROR1-Fc was performed using MabSelect SuRe (Cytiva), with PBS washing, elution using 50 mM acetic acid and neutralization with 10X PBS pH 7.4.

All purified proteins were analyzed by analytical size exclusion chromatography using an Agilent 1260 HPLC to assess native molecular weight and oligomeric state. Proteins were analyzed also for purity by reducing and non-reducing SDS-PAGE. Aliquots were flash frozen in liquid nitrogen and stored at -80°C.

#### Alexa Fluor 647 labeling

Proteins were thawed and adjusted to pH 8-9 with 1/10th volume of 20X Borate Buffer (Thermo Fisher Scientific), and incubated with Alexa Fluor 647 NHS ester (Thermo Fisher Scientific) overnight at 4°C with gentle mixing. Free dye was removed using either Zeba spin dye and biotin removal columns (Thermo Fisher Scientific) or purified by FPLC using a Superdex 200 prep grade HiLoad 16/600 column (Cytiva). Labeling was confirmed by measuring absorbance at 280 nm and 650 nm on a OneC Microvolume UV-Vis Spectrophotometer (Nanodrop).

#### Circular Dichroism

Protein secondary structure and thermal stability were measured by far-UV circular dichroism (CD) spectroscopy on a J-1500 spectrophotometer (Jasco). Protein samples at ∼0.25 mg/mL in PBS pH 7.2 buffer (Thermo Fisher) were analyzed in a 1 mm path-length cuvette. Sample temperature was maintained and adjusted using a Peltier-controlled cuvette holder. For wavelength scans to assess secondary structure, CD spectra were recorded from 300 to 195 nm at 25°C. In order to measure protein thermal stability, changes in CD absorbance at 208 nm were monitored as temperature was increased at 1°C/min from 25°C to 95°C.

#### Biolayer interferometry (BLI)

Binding of engineered cytokines to human IL-2Rα, IL-2Rβγ, and CD8α ligands was measured with BLI on an Octet RED96 system (ForteBio). All measurements were carried out at 25°C in a running buffer of HBS-EP+ (Cytiva) supplemented with 0.25% BSA. Ligands were first captured on suitable biosensors. After capturing ligands and establishing a baseline, biosensors were dipped into wells containing titrations of cytokine or VHH analyte to measure association and subsequently dipped into running buffer to measure dissociation. Specific binding conditions are described below.

To measure interactions between the designed cytokine and IL-2Rα, Fc-tagged IL-2Rα (R&D Biosystems) was captured on ProA biosensors (Sartorius) to ∼2 nm, and binding was measured with cytokine analytes serially diluted from 3 µM to 1 nM, using 360 s association and 60 s dissociation. To measure interactions between designed cytokines and CD8α, biotinylated CD8α (Acro Biosystems) was captured on streptavidin (SA) biosensors (Sartorius) to ∼0.6 nm, and binding was measured with analytes diluted from 100 to 0.4 nM, using 120 s association and 480 s dissociation. To measure interactions between designed cytokines and IL-2Rβγ, Fc-tagged IL-2Rβγ (Acro Biosystems) or biotinylated Fc-tagged IL-2Rβγ (Acro Biosystems) were captured on ProA or SA biosensors to ∼3 nm (SA biosensors were used for all VHH fusion proteins due to VHH binding to ProA); binding was measured with analyte serially diluted from 100 to 0.4 nM, using 120 s association and 480 s dissociation. All data was fitted using a 1:1 binding model in the Octet Analysis Studio software; affinities were calculated based on rates of association and dissociation (IL-2Rβγ and CD8α) or equilibrium binding responses (IL-2Rα). Measurements were performed in at least triplicate.

### *In vitro* functional assays

#### Cell isolation

Normal donor leukopaks (StemCell) were processed within 48h of receipt. Apheresis bags were mixed thoroughly and transferred to an appropriately sized conical. Cells were resuspended at a concentration of 50x10^6^ to 75x10^6^ cells/mL with EasySep Buffer (StemCell). A portion of the suspension was banked directly as frozen PBMCs. From the remaining suspension, total T cells were enriched by negative selection following the EasySep Human T Cell Isolation Kit (StemCell) and NK cells were enriched by negative selection following EasySep Human NK Cell Isolation Kit (StemCell). Selected T or NK cells were resuspended in CS10 (StemCell) at a concentration of 20x10^6^ or 10x10^6^ cells/ml, respectively, and frozen in either CoolCell containers (Corning) or in a controlled rate freezer (Thermo Fisher) according to manufacturer protocols, then transferred to long term cryostorage.

#### pSTAT5 assay

Leukopak-isolated PBMCs, T cells (resting T cells), T cells activated with 1% TransAct (Miltenyi) for 48h and expanded for 5 days before cryopreservation (activated T cells), or NK cells were thawed and rested overnight in the absence of exogenous cytokines, then mixed with T Cell Media (TCM; OpTmizer Basal Medium and Cell Supplement (ThermoFisher) with Immune Cell Serum Replacement (ThermoFisher), 2 mM L-glutamine (Gibco), and 2 mM Glutamax (Gibco)) containing the indicated recombinant cytokine and incubated for 15 minutes at 37°C. Cells were transferred onto ice for 5 min, washed 1x with cold Cell Staining Buffer (Biolegend), blocked for 15 min at 4°C with Human TruStain FcX (BioLegend) (NK cells and PBMCs only), and stained for surface markers for 20 min at 4°C. Cells were then fixed with Cytofix Fixation Buffer (BD) at 37°C for 10 min, washed, permeabilized with Phosflow Perm Buffer III (BD) for 30 min at 4°C, and stained for intracellular markers including pStat5 (pY694) (BD). The purified recombinant proteins used in these pSTAT5 assays are STII-tagged, with the exception of WT IL-15 (R&D Systems), which does not contain any tag.

#### *In vitro* proliferation assays

Total T cells or CAR-T cells were thawed and plated at 5e4 cells/well in TCM without exogenous cytokines. Where indicated, freshly thawed cells were stimulated for 24h with 1% TransAct, washed, and replated at 5e4 cells/well in TCM without exogenous cytokines. Every 3-4 days, cells were counted on a Cellaca PLX High-throughput Image Cytometer (Revvity) and split 1:2.

#### Vector production and titration

VSV-pseudotyped third generation lentiviral vector particles were generated using polyethylenimine (PEI)-mediated transient transfection of suspension-adapted 293Ts (ATCC) grown in shake plates in FreeStyle medium (ThermoFisher). Lentiviral supernatant was harvested, pooled, clarified, and filter-sterilized using 0.2µm filtration. Smaller batches of lentiviral supernatant were concentrated in 15mL conicals with Lenti-X PEG concentrator (3:1 v/v ratio). Supernatants are vortexed before incubating at 4℃ for 2 hours, followed by centrifugation at 3,000xg at 4℃ for 1.5 hours to pellet any vector bound to Lenti-X PEG. The volume was carefully aspirated and the pellet is resuspended to equal 10x concentration from the harvest volume. Larger batches of lentiviral supernatant were concentrated in 50 mL conicals for 4h at 4°C at 10,000 x g in 10% Sucrose Buffer ^40^. The volume was then decanted and pellets were resuspended to equal ∼60-100x concentration before vortexing vigorously. Concentrated lentivirus was aliquoted and stored at -80C. Lentiviral batch titers were evaluated by limited dilution on primary T cells and measured by flow cytometry against surface expression of the transduction marker and/or digital PCR (dPCR) against lentiviral Rev Response Element (RRE) using the following primer sets: Forward primer (5’-GCTTTGTTCCTTGGGTTCTTG-3’), Reverse primer (5’-TTCTGCTGCTGCACTATACC-3’), and Taqman probe (RRE-FAM: 5’-/56-FAM/AATTGTCTG/ZEN/GCCTGTACCGTCAGC/3IABkFQ/-3’). Digital PCR samples were analyzed on the QIAcuity Digital PCR system (Qiagen, Germantown, Maryland, USA). Functional titers ranged from 1.8 x 10^7^ - 1.7 x 10^8^ TU/mL. For *in vivo* experiments, commercial stocks of lentiviral vector were acquired from GenScript (GenScript) for three vectors, titers ranging from 1.6 x 10^8^ - 5.8 x 10^8^ TU/mL.

Lentiviral constructs were constructed using a 3rd generation backbone containing either a single promoter (MND or inducible) driving multiple cistrons separated by 2A ribosomal skip sequences, or two promoters arranged in reverse complement with an inducible promoter driving expression of the indicated cytokine, followed by a synthetic polyA site to prevent transcriptional readthrough into the constitutive MND promoter driving CAR and transduction marker. Ror1 targeting CAR was constructed with R12 ScFv, IgG4 spacer, CD28 transmembrane domain, 41BB costimulatory domain and CD3 zeta domain followed by a 2A ribosomal skip element and a truncated CD19 transduction marker downstream of the CAR ^41–43^. MSLN targeting CAR constructed with an anti-MSLN nanobody, IgG1 derived spacer, CD28 transmembrane domain, 41BB costimulatory domain and CD3 zeta domain, followed by a 2A ribosomal skip sequence and the EGFRopt transduction marker ^32^. Tested inducible promoters were composed of concatenated transcription factor binding sites and a minimal promoter.

#### Lentiviral transduction

For lentiviral transduction, T cells were cultured in TCM with IL-2 (200 IU/ml; R&D Systems), IL-7 (1200 IU/mL; R&D Systems), and IL-15 (200 IU/mL; R&D Systems). T cells were stimulated with 1% TransAct for 24h, followed by incubation with lentivirus. Stimulation and viral infection were terminated after 24h by addition of 7 volumes of fresh TCM with cytokines, without TransAct, and cells were expanded for 5 additional days before experimental use or cryopreservation in CS10 (StemCell).

#### Regulated Promoter design and screening

Promoters were designed using concatemers of known TF binding sites for transcription factors involved in T cell activation and cloned upstream of cytokine R2.1 to enable production upon T cell activation. Leukopak isolated T-cells were thawed and transduced with lentivirus and enriched by positive selection for the transduction marker CD19 following EasySep Human CD19 Positive Selection Kit II (StemCell) before flow cytometric validation 3 days post transduction. Cells were expanded for 11 days, then seeded at 0.5x10^6^ cells/mL in 1 mL TCM with no cytokines into either antigen-coated or uncoated plates. Antigen-coated plates were prepared by incubating 100 uL of 5 ug/mL of Ror1-Fc (in house) in PBS in a flat bottom 96 well plate the day of the assay, and washed 2x with sterile PBS prior to seeding cells. Cytokine production was quantified by STII MSD from supernatants collected 24 hours post stimulation.

#### TCR-T activation assay

T2 lymphoblasts (ATCC) were incubated with SLLQHLIGL peptide (Genscript) at the indicated concentrations for 1 hour, then washed with PBS (Gibco). TCR-T cells were incubated with 6.5 nM recombinant proteins for 15 min at 37°C, then washed twice. 2x10^5^ T2 cells and 1x10^5^ TCR-T cells, identified by expression of truncated EGFR (EGFRt), were cocultured in R10 (RPMI 1640 (Gibco) with 10% FBS (Gibco)) containing 1X transport inhibitor cocktail (Invitrogen). After 4 hours, cells were brought to single-cell suspension, stained for surface markers, fixed and permeabilized using Foxp3/Transcription Factor Staining Buffer Set (eBioscience), and stained for intracellular IFNg.

#### *In vitro* repeated stimulation assays

Freshly thawed CAR-T cells were rested overnight in TCM + IL-2, IL-7, and IL-15. CAR-T cells were enriched by positive selection for the transduction marker CD19 following EasySep Human CD19 Positive Selection Kit II (StemCell) before flow cytometric validation. 5x10^5^ CD19+ CAR-T cells were cocultured with 5x10^5^ H1975 tumor cells (ATCC) in a 12-well tissue culture plate. After 2 days, cells were stained with anti-human CD8 APC Viability Kit (Revvity) and anti-human CD19-PE (Biolegend) before analyzing on a Cellaca PLX High-throughput Image Cytometer (Revvity). Subsequently, 5x10^5^CD19+ CAR-T cells were transferred into a new 12-well plate containing 5x10^5^ H1975 tumor cells per well. This process was repeated for a third stimulation of 72h starting on day 4. On day 7, the frequency of CD19+ cells was determined by flow cytometry once more. 24h after each replating, 60 ul of supernatants were collected for cytokine analysis. All cocultures were performed in R10.

To stimulate MSLN CAR-T cells using immobilized MSLN/CD58/ICAM-1 proteins, wells of tissue culture plates were coated with approximately 155 ul/cm2 of 5 ug/ml of in-house manufactured recombinant human MSLN and 2 ug/ml each of recombinant human CD58-Fc and ICAM-1-Fc (R&D Systems). Plates were subsequently incubated at either 37°C for 2 hours or overnight at 4°C. After coating, residual coating solution was removed, and CAR-T cells diluted in TCM with cytokines were added to the wells. For acute stimulation, 1-2x10^6^ CAR-T cells/ml were incubated with immobilized MSLN/CD58/ICAM-1 for 24 hours. For chronic stimulation, 1-2x10^6^ CAR-T cells/ml were continuously cultured on MSLN/CD58/ICAM-1 coated wells for 9-10 days. To ensure continual access to stimulation proteins, CAR-T cells were enumerated and replated on freshly MSLN/CD58/ICAM-1 coated wells every 2-3 days (at 1-2x10^6^ CAR-T cells/ml) for a total of 4 rounds of stimulation over the 9-10 day period. After either acute or chronic stimulation, CAR-T cells are assessed in phenotypic or functional assays.

#### *In vitro* spheroid killing assay

Tumor spheroids were generated by adding 1e4 H1975 mKate2 or H1650 mKate2 cells to wells of a 96-well round-bottom ultra low attachment plate (Sbio) in 100 ul R10 media: RPMI 1640 + Glutamax-I (Gibco) with 10% FBS (Gibco). PBS (Gibco) was added to edge wells. The plate was centrifuged at 1,000xg for 10 min before transferring to a 37°C incubator for 72h.

T cells or freshly thawed NK cells were added to tumor spheroids at the indicated E:T ratios, in 100 ul total in R10 with the indicated recombinant cytokine, if any. Plates were imaged in an Incucyte Sx5 (Sartorius) every 6 hours for 7 days. Mean total red fluorescence integrated intensity (RCU x um2) was calculated for 3 technical replicates per sample and normalized to baseline for visualization.

#### NK cell proliferation assay

Upon thaw, NK cells were resuspended at 2x10^6^/mL in warm PBS and an equal volume of 2X concentration of CTV (ThermoFisher) in warm PBS was added to the cells for a final concentration of 2.5 um. Cells were incubated in a 37°C water bath for 9 minutes, protected from light, and then quenched with 5X volume of R10 media. Cells were centrifuged at 500xg for 3 min and recounted. CTV-labeled NK cells were then resuspended in media containing recombinant cytokines at 2X the indicated final concentration. NK samples were then added to tumor spheroids or plain media at a 2:1 E:T ratio in 100 uL. Samples were placed in a 37°C incubator for one week. On day 4, half the media was removed from each well and replaced with fresh media containing the indicated recombinant cytokine, if any.

After one week, samples were centrifuged at 1500xg for 30 seconds and resuspended in 100 uL CSB (Biolegend). Technical duplicates were combined, and after letting remaining spheroids settle to the bottom of the wells (30 seconds), 100 uL of each sample was removed off the top and transferred to a new 96 well plate for staining. Cells were stained for surface markers for 20 min at 4°C, and then fixed with Fluorofix Buffer (BD) at 4°C for 20 min. Precision Count Beads (Biolegend) were included in the surface stain to enable calculation of absolute cell number.

#### Endogenous cytokine MSD

Samples collected from in vivo and in vitro assays were assessed for IFNg and IL-2 expression using V-PLEX Proinflammatory Panel 1 Human Kit (MSD K15049D-4) following manufacturer’s instructions with the following changes. Calibrators were 4X more concentrated than recommended to keep samples on scale. Additionally, sample and detection antibody were added simultaneously. NA values were imputed with 0.

#### STII MSD

Strep-tag II (STII)-tagged protein concentrations were measured using a custom MSD assay. Standard 96-well MSD plates (MSD L15XA-3) were coated overnight at 4°C with 30 µL/well of 5 µg/mL polyclonal anti-human IL-2 capture antibody (BioLegend custom order Poly5176 900005704) in PBS. After incubation, plates were washed three times with 1X PBST (Thermo-Fisher). Recombinant IL-2-STII protein in RPMI + 10% FBS was used to prepare the standard curve ranging from 50000-0.64 pg/mL. Each well received 25 µL of sample (protein standard or unknown) and 25 µL of anti-STII detection antibody. The anti-STII detection antibody (GenScript) was labeled in-house with SULFO-TAG NHS-Ester (MSD A01732) according to manufacturer’s instructions and added at 0.5 µg/mL in Diluent 100 (MSD R50AA-2). After shaking at 700 RPM for 2h at room temperature, plates were washed 3 times with 1X PBST and 150 µL/well of 2x Read Buffer T (MSD R92TC-1) was added. Plates were read on an MESO QuickPlex SQ 120MM and analyzed with the MSD Discovery Workbench software. NA values were imputed with 0.

### *In vivo* experiments

#### *In vivo* tumor efficacy

The H1975 ROR1 CAR-T tumor efficacy study was carried out under contract at CRL (Durham, NC) using NOD-*Prkdc^em26Cd52^Il2rg^em26Cd22^*/NjuCrl (NCG) mice (Charles River) bearing tumors implanted subcutaneously at 10x10^6^ cell/mouse. The H1975 tumor cells were resuspended in 100uL of PBS and mixed 1:1 with 100uL of Matrigel (Corning) for a 200uL injection in the right flank. After implant, mice were monitored twice weekly for body weight and tumor volume performed using caliper measurements and the formula (width^2^ x length)/2 for calculating tumor volume. Eight days post implant, mice were randomized into study groups of 5 with mean volumes falling between 98mm^3^-102mm^3^. Frozen vials of engineered CAR cells from two different human donors were thawed and formulated to contain either 1x10^6^ or 2x10^6^ total CAR^+^ cells and injected in 200uL of PBS. Mock cells were dosed to match the maximum total cell number of the highest dose group for each donor. Injections were administered intravenously via tail vein. Animals exited study when either a maximum tumor burden of 2000mm^3^ was reached or when body weight dropped below 70% of baseline, whichever occurred first. Peripheral blood was collected 6 days post CAR injection via the submental route (100uL) into EDTA coated tubes (BD) and processed for flow cytometry. For staining, a set volume of blood was lysed in ACK buffer (Quality Biological, Inc.), neutralized with PBS, and stained using mCD45-BV650, hCD45-PE-Cy7, CD3-FITC, CD4-BV785, CD8-BV605, CD39-APC, CD19-BV421, PD-1-PE (BioLegend) and a Live/Dead dye (Thermo-Fisher). Samples were acquired on a LSR-Fortessa (BD) and analyzed using FlowJo software (Tree Star, Inc; v10.0.7r2). Study was performed under approved IACUC protocols, and in compliance with all local and federal regulations.

The H1650 MSLN CAR-T tumor efficacy study was carried out by Outpace Bio staff at a CRL Accelerator and Development Lab (CRADL) in Seattle, WA using NOD.Cg-*Prkdc^scid^Il2rg^tm1Wjl^*/SzJ (NSG) mice (Jackson Laboratory) implanted subcutaneously with 5x10^6^ cells/mouse. The H1650 cells were resuspended in 100uL of PBS and mixed 1:1 with 100uL of Matrigel (Corning) for a 200uL injection in the right flank. Monitoring initiated as outlined above. Twenty days post implant when tumor volumes averaged 164.9mm^3^, mice were randomized into study groups of 5 and dosed with the corresponding number of engineered CAR-T cells contained within 5x10^6^ total cells via intravenous tail vein injection. Animals exited study when either a maximum tumor burden of 2000mm^3^ was reached or when body weight dropped below 80% of baseline, whichever occurred first. Peripheral blood (200uL) was collected as described above and processed for flow cytometry. For plasma isolation, 100uL of blood was mixed with 100uL of PBS and spun at 1000xG for 5 minutes after which 100uL clarified plasma was removed and frozen at -80°C for future cytokine analysis. Blood samples were stained using 50uL of whole blood in a total of 100uL with CD45RA-BUV395 (BD), CD4-BUV496 (BD), CD62L-BUV615 (Thermo Fisher), EGFR-BV421, CD39-BV510, CD3-BV650, CD8-BV785, TIGIT-PE, hCD45-PE-Dazzle594, and PD-1-PE-Cy7 (Thermo Fisher); all antibodies were BioLegend unless otherwise stated. Precision counting beads (BioLegend) were included for calculating cells/mL and after staining samples were lysed with RBC lyse/fix solution (BioLegend) and acquired on a ZE5 (BioRad) and analyzed using FlowJo software (Tree Star, Inc). Tumor samples were collected at end of study, weighed, and dissociated into a single cell suspension using the tumor dissociation kit and GentleMACS instrument running the 37C_h_TDK-2 protocol (Miltenyi Biotec). Tumor samples were then stained using the above protocol, processed, and analyzed in the same manner. Study was performed under approved IACUC protocol EB17-010-407 and in compliance with all local and federal regulations.

#### Statistical analysis

All *in vivo* tumor killing analyses were performed by fitting linear models to log-transformed tumor volume AUC with log-transformed baseline tumor volume and treatment group (coded as 0 for reference group and 1 for comparison group). For any two treatment groups to be compared, AUC was computed via the trapezoid rule over the period spanning from administration of treatment to the latest study time point at which at least 3 mice are present in each group.

Missingness due to mouse dropout was addressed via multiple imputation. Within each group, imputations are performed using the pan() function in the pan R package via a model including fixed effects in day (categorical) as well as random intercepts and slopes in day (continuous, centered) for each mouse. 5 sets of imputations are produced for each mouse with missing data. For each imputed dataset, AUC was computed for each mouse and the linear model described above was fit. Estimates and standard errors from these models are combined using the pool() function in the mice R package. Wald p-values for a null hypothesis of no treatment effect are reported.

Evidence for differences in mouse survival curves across treatment groups (figure 3h) was evaluated via log-rank tests as implemented in the survdiff function in R package survival. Possible differences in mean relative cell count (figure 3i) are evaluated via robust Poisson regression of these counts on treatment group (coded 0 for reference group and 1 for comparison group) using the glmtest function in the raoBust; p-values from robust score tests of no difference in mean relative cell count are reported. Evidence for differences in mean percent CD8+ T cells was evaluated via Welch’s t-test.

Within in vitro experiments, evidence for differences across cytokine treatment group in geometric mean (GM) NK cell count as well as in GM relative endpoint fluorescence (figures 4d and e) are evaluated via paired t-tests applied to log-transformed measured values, with pairing on T cell donor.

All analyses were performed using R version 4.5.0.

- R Core Team (2025). R: A Language and Environment for Statistical Computing. R Foundation for Statistical Computing, Vienna, Austria. <https://www.R-project.org/>
- Stef van Buuren, Karin Groothuis-Oudshoorn (2011). mice: Multivariate Imputation by Chained Equations in R. Journal of Statistical Software, 45(3), 1-67. DOI 10.18637/jss.v045.i03
- Jing Hua Zhao and Joseph L. Schafer (2023). pan: Multiple imputation for multivariate panel or clustered data. R package version 1.9.
- Willis A, Clausen D, Teichman S, Mathur S (2025). raoBust: Robust Score Tests for Poisson and Multinomial Regression. R package version 0.1.3.3. <https://github.com/statdivlab/raoBust>
- Therneau T (2024). A Package for Survival Analysis in R. R package version 3.8-3, <https://CRAN.R-project.org/package=survival>

